# The origins and adaptive consequences of polyploidy in a dominant prairie grass

**DOI:** 10.1101/2025.11.25.690567

**Authors:** Alyssa R. Phillips, Taylor AuBuchon-Elder, Kerrie Barry, Michelle Stitzer, Edward S. Buckler, Robert Bukowski, Brenda Cameron, Elli Cryan, Elisabeth Forrestel, Paul P. Grabowski, Jane Grimwood, Jerry Jenkins, Shengqiang Shu, Anna M. Lipzen, John T. Lovell, Patrick Minx, Julianna Porter, Daniel Runcie, Jeremy Schmutz, Britney Solomon, Jessica Stephens, Qi Sun, Melissa Williams, Yuko Yoshinaga, Sherry Flint-Garcia, M. Cinta Romay, Elizabeth A. Kellogg, Jeffrey Ross-Ibarra

## Abstract

Polyploidy is ubiquitous across North American prairies, which provide essential ecosystem services and rich soil for agriculture. Yet the mechanism driving polyploid abundance is unclear. Multiple hypotheses have been proposed including polyploid abundance is proportional to the opportunity for whole genome duplication (WGD), and WGD alters phenotypes that may increase fitness. We tested these two hypotheses together in the mixed-ploidy species *Andropogon gerardi*, a dominant grass species in endangered North American tallgrass prairies. Leveraging a novel, phased allopolyploid reference genome, we found the *A. gerardi* hexaploid arose after the C_4_ grassland expansion in the early Pleistocene, when glacial cycles likely increased secondary contact between the diploid progenitors. We sequenced *A. gerardi* from 25 populations and examined cytotype performance and morphology in a controlled environment to investigate the consequences of the contemporary mixed-ploidy populations. We found the 9*x A. gerardi* cytotype is a neopolyploid and a result of recurrent WGD events. Further, we demonstrate the 9*x* neopolyploids have greater growth and a decreased stomatal pore index, which is adaptive in xeric climates where the 9*x* cytotype is most common. Together, our results support both hypotheses for polyploid abundance in North America: WGD is a product of opportunity and can have immediate fitness consequences. Although the changes to fitness may provide an advantage to 9*x A. gerardi*, the establishment of 9*x* may lower overall population fitness due to the lower reproductive viability of 9*x* individuals.

**Significance statement:** Polyploid species are abundant in North American prairies and make up many of the dominant species in the ecosystem. This prominence could be a result of whole genome duplication conferring an advantage that increases the frequency of polyploids or could simply indicate that the opportunity for whole genome duplication is higher in this ecosystem, or both. Through examining three polyploidization events in *A. gerardi*, the dominant species in endangered tallgrass North American prairies, we found whole genome duplication is both surprisingly common and confers traits that are beneficial in some environments.

## Introduction

Polyploid grasses have dominated North American grasslands since at least the C_4_ grassland expansion in the Late Miocene, a period of dramatic global change (Estep *et al*. 2014; Edwards *et al*. 2010). These include economically important taxa (*Panicum virgatum* and *Zea* spp.) as well as ecologically dominant species (*Andropogon gerardi*, *Schizachyrium scoparium*, *Sorghastrum nutans*, *Bouteloua gracilis*) which are essential for ecosystem function (Stebbins 1975; Avolio *et al*. 2019). These grasses represent a diverse continuum of polyploidy, from allopolyploids which form through hybridization of two or more species, to autopolyploids formed through genome doubling within a single species (Stitzer *et al*. 2025; Estep *et al*. 2014), and many exhibit variation in polyploidy within a species (Stitzer *et al*. 2025; Delaney and Baack 2012).

The prevalence of polyploidy in this ecosystem has confounded scientists, as new polyploids are not favored to survive. The establishment of new polyploids is challenged by a demographic bottleneck and a frequency-dependent mating disadvantage that arises when the only mates available are the diploid progenitor (minority cytotype exclusion, Levin 1975). Further, new polyploids must overcome challenges associated with higher DNA content, increased resource requirements, meiotic abnormalities, epigenetic instability, and altered gene dosage and allele number (Bird *et al*. 2018; Ramsey and Schemske 2002). Together, these challenges make the establishment of polyploidy from a small number of founders relatively unlikely (Oswald and Nuismer 2011).

Multiple hypotheses have been proposed for how polyploids overcame these challenges and rose to their current prominence. Stebbins (1985) proposed polyploid abundance is related to the opportunity for poly-ploidization. Paleoclimatic events may have altered migration patterns and increased the probability of secondary contact between species with limited reproductive barriers, thereby increasing the probability of allopolyploidy (Stebbins 1985). The Pleistocene glaciations may have served as this event in North America, forcing secondary contact in limited habitable space, and resulted in high numbers of polyploids which could overcome minority cytotype exclusion and the initial demographic bottleneck (Stebbins 1975). Polyploid frequency could also be determined by variation in the rate of unreduced gamete formation and thus the probability of polyploid embryos forming. Unreduced gamete formation is estimated to occur at a rate of 0.5-2.5% and given that many of these species may produce millions of gametes each year, polyploidization could be common (Kreiner *et al*. 2017; Ramsey and Schemske 1998). Consistent with this idea, an increasing number of polyploid species have been shown to have multiple independent origins rather than a single origin (Napier *et al*. 2022; Husband and Sabara 2004; Parisod and Besnard 2007; Ramsey *et al*. 2008; Servick *et al*. 2015; Monnahan *et al*. 2019; Edger *et al*. 2025). Alternatively, polyploid abundance could be driven by novel adaptive variation generated through WGD, enabling polyploids to outcompete their diploid progenitors.

Polyploidy has been shown to increase cell and plant size and alter floral morphology (Clo and Kolář 2021; Porturas *et al*. 2019). Although these effects are not always consistent across taxa, they can result in assortative mating between cytotypes and may aid survival in new or changing environments (Van de Peer *et al*. 2021). Estep *et al*. (2014) estimated a high frequency of polyploid speciation in Andropogoneae, a globally distributed clade that dominates tallgrass prairies at the beginning of the C_4_ grassland expansion in the Late Miocene and proposed that adaptive advantages of polyploidy may have enabled the expansion. Both of these hypotheses are likely tied to environmental change: the environment may effect the opportunity for polyploidization as unreduced gamete formation can be elevated under environmental stress (Wang *et al*. 2017; Sax 1936; Belling 1925), and the environment may select for or against novel polyploid phenotypic variation. Linking the processes that govern polyploid formation and establishment is necessary to understand the role of polyploidy in North American grasslands.

The dominant species in endangered North American tallgrass prairies, *Andropogon gerardi* Vitman (formerly *A. gerardii*; within Andropogoneae), serves as a particularly useful system for investigating the abundance of polyploids. *A. gerardi* is a mixed-ploidy species composed of hexaploids (2*n* = 6*x* = 60) and enneaploids (2*n* = 9*x* = 90) that live in sympatry (Keeler *et al*. 1987; Norrmann *et al*. 1997). The 6*x* cytotype is an allopolyploid estimated to have arisen between 3-9 Mya (Stitzer *et al*. 2025; Estep *et al*. 2014) during a period of dramatic global climate change, but the order of polyploid formation is unclear. Whereas the 9*x* cytotype is an autopolyploid of a reduced and unreduced 6*x* gamete (Norrmann *et al*. 1997). The two primary cytotypes of *A. gerardi* are prevalent range-wide and commonly found in sympatry (Tompkins *et al*. 2015; Keeler *et al*. 1987; Keeler 2004; Norrmann *et al*. 1997; Keeler 1990) while other cytotypes are rare, occurring in less than 5% of samples (McAllister *et al*. 2015; Keeler 1992). Contemporary North American tallgrass prairies are highly endangered, having experienced widespread decline from land-use change, and further decline is expected with climate change (Weaver 1968; Samson and Knopf 1994; Smith *et al*. 2017).

Mixed-ploidy populations of *A. gerardi* enable study of the effects of polyploidy that are not confounded by evolutionary differences between species. McAllister and Miller (2016) found at least 3 origins of the 9*x* cytotype by using reduced representation sequencing. It remains unclear, however, whether the widespread distribution of the 9*x* cytotype stems from a few WGD event followed by range expansion or from many local WGD events. Previous cytological and experimental research supports the latter hypothesis as the 9*x* cytotype exhibits low rates of successful sexual reproduction; the odd-ploidy of the 9*x* cytotype creates meiotic errors which results in aneuploid offspring with reduced life span in both inter-and intraploidy crosses (Norrmann *et al*. 1997; Tompkins *et al*. 2015). Therefore, 9*x* range expansion by sexual reproduction from only three origins is unlikely. Rather, the 9*x* cytotype may have expanded clonally via an underground rhizome, which is a common growth mechanism generating as much as 99% of the recruitment of new shoots in some environments (Benson and Hartnett 2006).

The environment may also play a role in the abundance of the 9*x* cytotype. Although both cytotypes are found across the range of *A. gerardi*, the 9*x* cytotype has higher abundance in drier climates with higher diurnal and annual temperature ranges, such as the southwestern United States (McAllister *et al*. 2015). The correlation of 9*x* abundance with climate suggests that coexistence, establishment, or survival of the polyploid cytotype may also depend on the environment. For example, two experiments comparing seed set of 6*x* and 9*x* found conflicting results between *A. gerardi* populations; the 9*x* plants either produced a comparable amount of viable seed to the 6*x* (Colorado and Nebraska) (Keeler and Davis 1999) or very low amounts of viable seed (North Carolina) (Tompkins *et al*. 2015), although viable seed is expected to be aneuploid. Previous work has identified ecotypic variation in *A. gerardi* that follows the same environmental clines as 9*x* abundance. The described ecotypes vary in leaf anatomy, plant height, and physiology (Olsen *et al*. 2013; Caudle *et al*. 2014; Galliart *et al*. 2020; Bachle and Nippert 2021), but trait variation has been examined only in a limited part of the range and without consideration of polyploidy.

Here, we investigate the evolution of polyploidy in *A. gerardi* and the consequences of WGD on phenotype and adaptation. We assess whether polyploidy is a product of opportunity by leveraging the first chromosome-scale assembly and a range-wide sample of *A. gerardi* genomes to describe the origins of 6*x* and 9*x* cytotypes. Then, we evaluate whether WGD conveys adaptive phenotypic variation using performance and leaf economic traits measured in a two-year common garden experiment with genotypes from 14 populations. Combined, our evidence for repeated WGD and phenotypic evaluation of ploidy differences suggest polyploid establishment and abundance is a product of multiple mechanisms working in concert.

## Results

### Allopolyploid formation of *A. gerardi* occurred in two steps

To facilitate our evolutionary analyses, we assembled the first haplotype-resolved A. gerardi reference genome (accession Kellogg-1272). Using a combination of 92x coverage Illumina short-reads, 54x OmniC reads, and 85x PacBio HiFi long reads (Table S1), we assembled 30 chromosome-scale scaffolds per haplotype representing 99.9% of the HAP1 and HAP2 assemblies (Table S5-S6). The assembly is highly contiguous and complete, with both haplotypes having a scaffold N50 of 13 Mb (Table S5-S6) and 0.001% gaps in HAP1 and 0.05% gaps in HAP2. We generated gene models and repeat-content annotations for both haplotypes. Approximately 70% of HAP1 and HAP2 was annotated as repetitive elements, and contained 89,426 and 88,437 gene models, respectively, consistent with other grass genomes (Hufford *et al*. 2021; Wicker *et al*. 2018; Stitzer *et al*. 2025). Gene space completeness was measured using BUSCO (Manni *et al*. 2021). The assembly contains 99.7% (HAP1; HAP2 is 98.7%) of the expected conserved genes, 59.7% of which are duplicated.

Previous work suggests that the diploid and tetraploid parental species likely belong to *Andropogon* (recently split into *Andropogon* and *Anatherum*, Vorontsova *et al*. 2023) and *Schizachyrium* (Estep *et al*. 2014; Nagahama and Norrmann 2012). We leveraged this information to resolve the three subgenomes within the reference. We initially separated the chromosome-scale scaffolds into the three subgenomes (Fig. 1D) based on shared enriched k-mer content. Then, we used a publicly available reference genome for *Anatherum virginicum* (formerly *Andropogon virginicus*, Vorontsova *et al*. 2023) to assess which subgenome is most closely related to *Andropogon*. The subgenome that shared the highest number of 18-mers with *A. virginicum* was labeled subgenome’A’.

**Figure 1:**
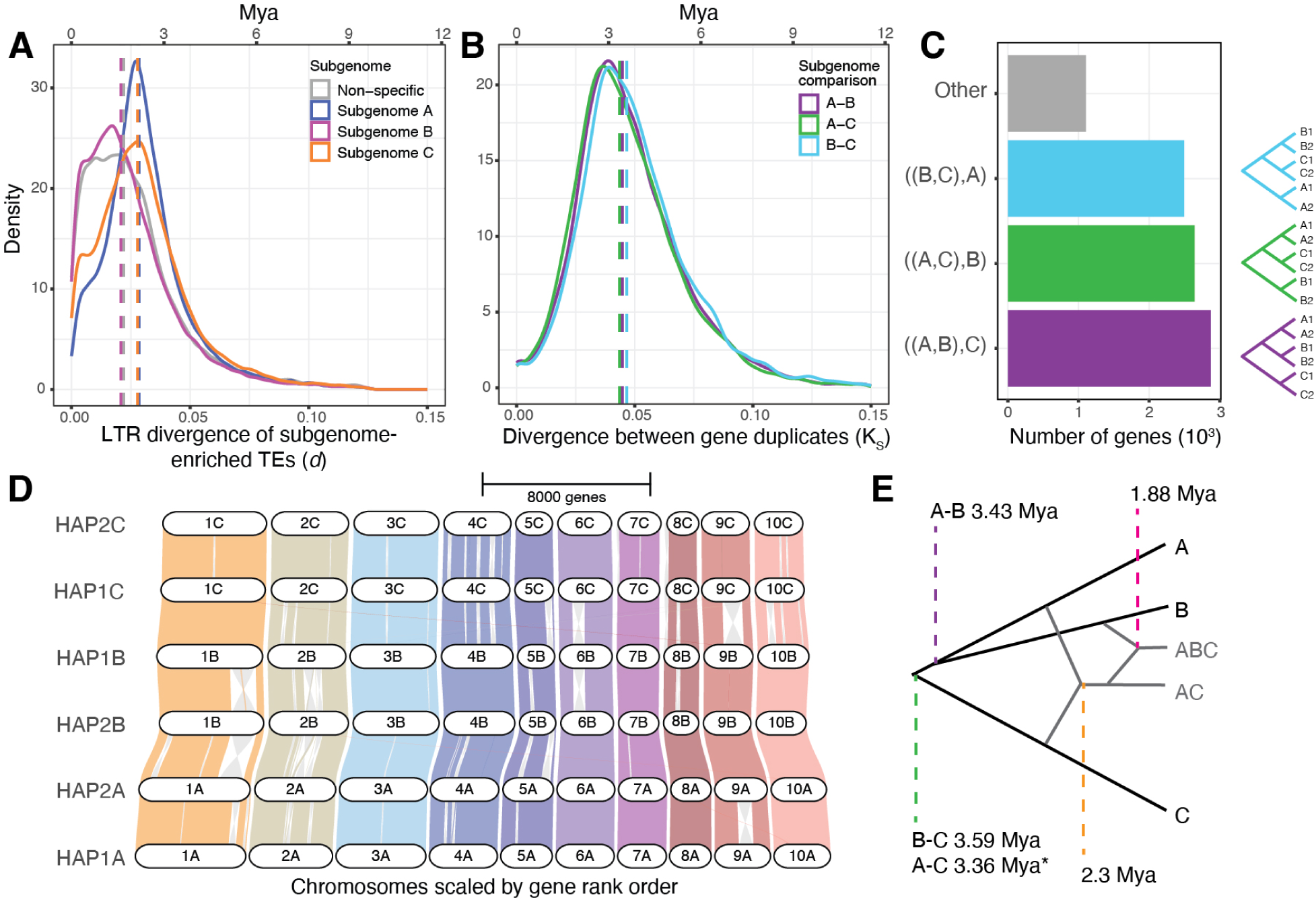
T**h**e **evolutionary history of the *A. gerardi* subgenomes. (A)** LTR divergence (*d*) of subgenome-enriched TEs within each subgenome, with the median LTR divergence of each subgenome plotted as a vertical line. **(B)** Synonymous site divergence (*K_S_*) of syntenic gene duplicates in A. gerardi, with the mean *K_S_* plotted as a vertical line. **(C)** Counts of gene tree topologies of 9,111 syntenic loci. **(D)** Syntenic map of *A. gerardi* subgenomes and haplotypes estimated with GENESPACE (Lovell *et al*. 2022). The colored ribbons between chromosomes indicate the orthologous regions among genomes, with gray ribbons indicating inversions. **(E)** Graphic of the proposed model of *A. gerardi* hexaploid formation with node dates from A and B. Asterisk indicates possible gene conversion leading to underestimation of divergence time.

Leveraging our phased reference genome, we examined the timing and order of the polyploidization events forming the allohexaploids. We used the biology of transposable elements (TEs) to estimate the timing of the duplication events giving rise to modern *A. gerardi*. TE content differs among even closely related species (Lee and Kim 2014), but after two genomes are combined into a single nucleus, new insertions occur homogeneously across the subgenomes (Session and Rokhsar 2023), making TEs useful markers to track lineages within allopolyploids. The age of long terminal repeat (LTR) retrotransposons can be estimated from differences between their flanking repeats, allowing us to identify the timing of polyploid formation and cessation of subgenome-specific TE amplification. We employed Subphaser (Jia *et al*. 2022) to identify 13,482 LTR retrotransposons enriched in subgenome A and 16,292 and 12,541 in subgenomes B and C, respectively.

A Kruskal-Wallis test showed significant differences between the timing of insertion among subgenomes. We found subgenomes A and C show similar median timings of insertion (Fig. 1A; A *d* = 0.0297, 2.28 Mya; C *d* = 0.0301, 2.32 Mya; post-hoc Dunn’s test not significantly different p = 0.7383), while subgenome B insertions are younger (B *d* = 0.0244, 1.88 Mya; significantly different from both A and C, p *<* 0.001). These results suggest an initial tetraploidy of A-C, followed by hexaploidy with the contribution of subgenome B. Previous cytological and genetic studies suggested three parental lineages of the hexaploids, with two being closely related (Estep *et al*. 2014; Nagahama and Norrmann 2012). To understand the phylogenetic relationship among the three parents, we measured pairwise divergence at synonymous sites and estimated gene tree topologies across 9,111 syntenic genes. Genic pairwise differentiation was significantly different between all subgenome comparisons (Kruskal-Wallis test, *χ*^2^ = 88.19, df = 2, p *<* 2.2e-16; post-hoc Dunn’s test, p *<* 0.001): the subgenome A-C comparisons showed the lowest median differentiation (A-C *K_S_* = 0.0437, 3.36 Mya), A-B intermediate (A-B *K_S_* = 0.0446; 3.43 Mya), and B-C the greatest (B-C *K_S_* = 0.0467; 3.59 Mya; Fig. 1B). Gene conversion and recombination may have homogenized the A and C subgenomes after the initial tetraploidy, perhaps explaining why A and C had the lowest differentiation. The B subgenome comparisons suggest it is more closely related to A than to C. In accordance with this, gene tree topologies for the 9,111 syntenic loci recovered similar proportions of sister relationships (A-B 31.5%, A-C 29.0%, B-C 27.4%; Fig. 1C), but placed the A and B subgenomes as sister taxa. The vast majority of gene trees placed the haplotypes of each subgenome together (87.9%), consistent with allopolyploidy and cytological studies reporting limited multivalent formation (Norrmann and Keeler 2003). Under this model, the A and B subgenomes form a sister pair, with gene conversion following polyploidy homogenizing differences between subgenomes A and C (Fig. 1E). While we cannot rule out an alternative topology in which the A and C diploid progenitors form a clade, perhaps with differential gene flow between taxa, our data support the formation of the current *A. gerardi* hexaploid cytotype by 1.88 Mya from three progenitor diploid lineages that diverged approximately 3.5 Mya (likely from *Andropogon* and *Schizachyrium*; Nagahama and Norrmann 2012).

### *A. gerardi* populations exhibit limited population structure and high genetic diversity

Although many *A. gerardi* are allohexaploids, cytological and flow cytometry analyses have shown that enneaploid (9*x*) individuals — with a third copy of each of the three subgenomes — are common and are found intermixed with allohexaploids (6*x*) in many different populations (Keeler *et al*. 1987; Norrmann *et al*. 1997; McAllister *et al*. 2015; Tompkins *et al*. 2015; Keeler 1990, 2004). To study the evolution of mixed-ploidy in *A. gerardi*, we assembled a diverse panel of 180 plants from 25 populations, representing the most geographically and environmentally dispersed WGS sample of *A. gerardi* to date (Fig. 2A). The majority of samples were sequenced to low coverage (*<*5x) and a subset was sequenced to high coverage (*>*20x) for quality control. We aligned reads to HAP1 of our *A. gerardi* assembly and, to correct for biases due to mixed-ploidy, based our subsequent analyses on single-read genotypes called for 100,000 random sites (see Methods).

**Figure 2:**
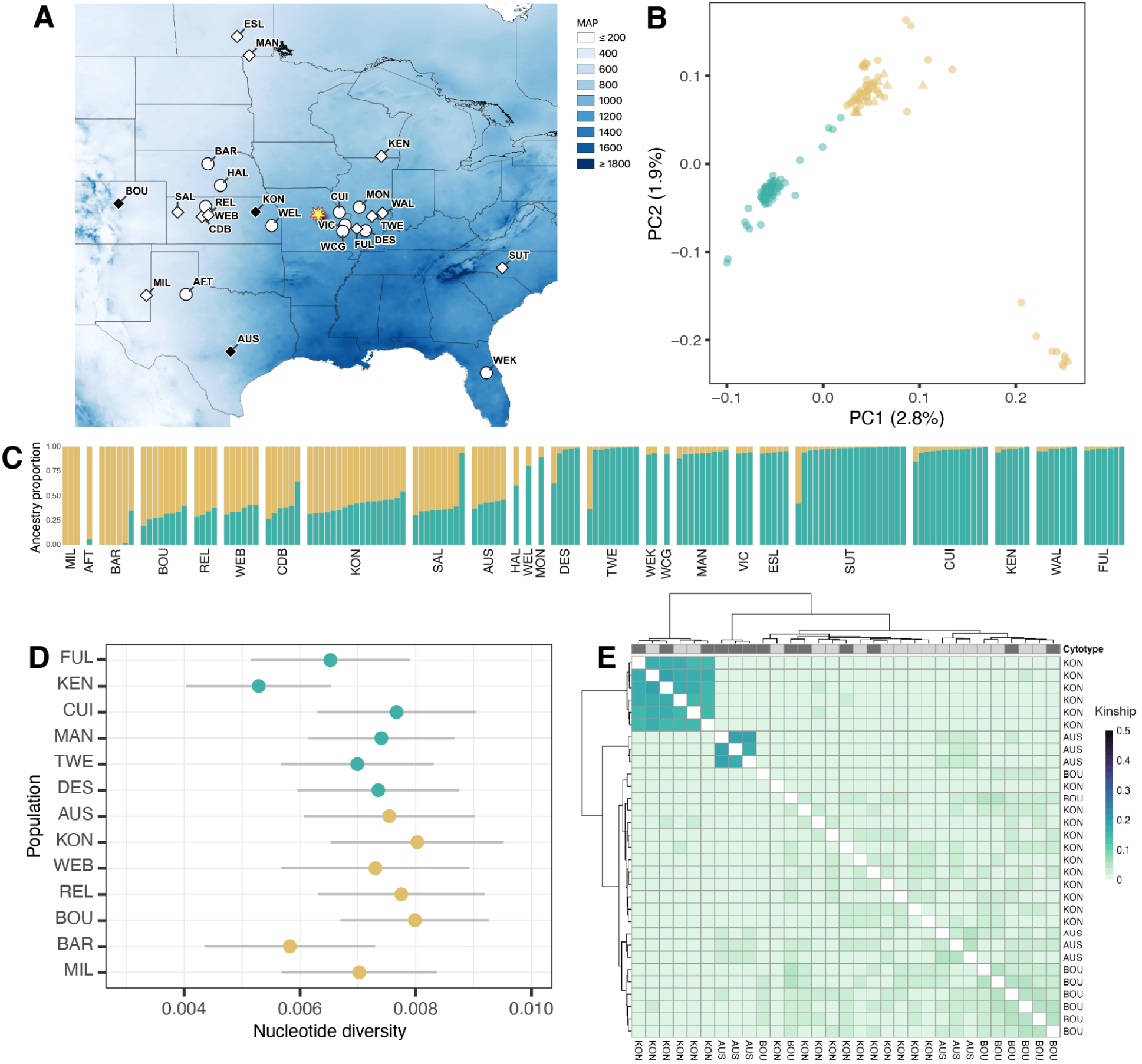
P**o**pulation **structure is limited among sampled *A. gerardi* populations. (A)** The geographic distribution of sampled populations relative to the gradient of mean annual precipitation across North America. Populations where only 6*x* genotypes were sampled are depicted in white and populations with at least two 9*x* genotypes and 6*x* sampled genotypes are shown in black. Diamond-shaped points indicate populations that were included in the common garden. Population codes refer to those in Dataset 01. The common garden site is indicated with a yellow star, and a photo of the garden is inset. **(B)** The first two PCs from a PCA on 100,000 random SNPs for all genotypes. Points are colored by the majority ancestry from Panel C. **(C)** Plot of STRUCTURE results when K=2 showing the ancestry proportion estimated for each genotype. The 9*x* genotypes are marked with a triangle. **(D)** Nucleotide diversity estimated per population for 6*x* genotypes. Populations are sorted by mean proportion of ancestry to the West admixture group. **(E)** Kinship among genotypes from mixed-ploidy populations subset from a larger kinship matrix of the West admixture group (Fig. S6). Genotypes are labeled by population (AUS, BOU, or KON). Rows and columns are hierarchically clustered by Euclidean distance in kinship vectors. Columns are annotated with the cytotype of each genotype where 9*x* are dark gray and 6*x* are light gray.

The 9*x* and 6*x* cytotypes of *A. gerardi* are nearly indistinguishable in the field, but flow cytometry and sequencing data enabled us to identify 9*x* individuals in six sampled populations, concentrated in the western half of the species range (Dataset 01), consistent with previous reports of increased 9*x* abundance in this region (McAllister *et al*. 2015). Three of the six populations contained multiple 9*x* and 6*x* individuals (BOU, KON, AUS; Fig. 2A).

We assessed overall levels of population structure in the sampled *A. gerardi* populations and how the two cytotypes are associated with this structure. Genetic diversity across the 6*x* cytotypes is high (mean *θ_π_* across populations = 0.007, Fig. S1), comparable to other perennial outcrossing taxa (Evans *et al*. 2018), and does not reflect the recent extensive habitat fragmentation of North American prairies (Samson and Knopf 1994). Notably, nucleotide diversity within *A. gerardi* is substantially less than *K_S_* between subgenomes, supporting that subgenomes reflect an allopolyploid, rather than autopolyploid, origin. A principal component analysis (PCA) of all samples, including both 6*x* and 9*x*, reveals minimal evidence of population structure. Although the first two principal components (PCs) broadly separate populations from the eastern and western portions of the range (referred to as the East and West group; Fig. 2B, S2), combined they explain less than 5% of the genetic variation across all samples. Consistent with this, differentiation between the East and West genetic groups was very low (*F_ST_* = 0.023±0.04; *ρ* = 0.043±0.068). Model-based admixture analysis further supports the East-West distinction (Fig. 2C, S3) and these genetic groups are consistent with previous genotyping studies (McAllister and Miller 2016; Galliart *et al*. 2020; Gray *et al*. 2014). Together PC1 and PC2 also distinguish a subset of genotypes within the West group; we estimated these genotypes to have high estimated relatedness (belonging to MIL and AFT; Fig. S6) and significantly higher average inbreeding coefficients (belonging to BAR, Fig. S7), thus the PCA revealed this pattern of high relatedness. STRUCTURE run without these three populations resolved the same East and West groupings (Fig. S4). Further, a PCA run on the high-coverage data subset with genotype likelihoods estimated in ANGSD (Korneliussen *et al*. 2014) demonstrated similar structure (Fig. S5). Neither the PCA nor the admixture analysis separated the 9*x* and 6*x* cytotypes (Fig. 2B-C), consistent with previous work suggesting there is no single range-wide origin of the 9*x* cytotype (McAllister and Miller 2016). However, these analyses did not provide the resolution needed to examine more detailed intraspecific relationships.

### Mixed-ploidy is maintained by recurrent polyploidization

To better understand the multiple origins of the 9*x* cytotype at a finer scale, we assessed whether the 9*x* cytotype arose from just a few origins followed by clonal expansion or many independent WGD events. If the 9*x* genotypes from different populations arose through a shared WGD event, the genetic similarity among 9*x* genotypes would be higher than similarity among 6*x* genotypes from the same populations due to a bottleneck associated with the WGD event. We estimated *ρ*, a genetic differentiation statistic comparable to *F_ST_* which was developed to overcome bias introduced by polyploidy (Ronfort *et al*. 1998), between the three sampled mixed-ploidy populations in the West genetic group (AUS, BOU, KON). Genetic differentiation among 6*x* from the mixed-ploidy populations 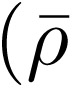 = 0.062) was not significantly different than differentiation among 9*x* 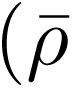 = 0.090; t-test t =-5.6242, df = 2.0848, p = 0.02748; Fig. S8, S9), largely rejecting the possibility of a single origin of the 9*x* cytotype in the West. To evaluate the role of clonal growth in maintenance of the 9*x* cytotype, we calculated kinship (*F_ij_*) between all 6*x* and 9*x* genotypes in the West genetic group. The majority of sampled genotypes had very low relatedness 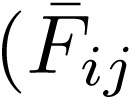 = 0.0078, SD = 0.021; Fig. 2E, S6), with the average kinship among 9*x* estimated as 0.017 (SD = 0.037 with a max of 0.18), clearly inconsistent with clonality. Together, these findings suggest mixed-ploidy in *A. gerardi* populations is a product of recurrent polyploidization events within individual populations.

### Polyploidy affects growth rate and stomatal traits

Whole genome duplication is well known to affect a number of plant phenotypes that may impact fitness in different environments (Clo and Kolář 2021). We conducted a 2-year common garden experiment of individuals from across our sequenced populations (Fig. 3A) to assess whether the excess of 9*x* genotypes in regions with high temperature variability and low mean annual precipitation is related to the phenotypic impacts of polyploidy. Using linear mixed models, we evaluated the effect of cytotype on 15 ecologically relevant phenotypes in 84 genotypes from 14 populations, including three populations (BOU, KON, AUS) with multiple 9*x* and 6*x* genotypes (Dataset 01). The measured phenotypes (Fig. 3, SI Appendix) were selected to provide estimates of leaf economics, stomatal traits, and performance (reproductive effort and growth) that describe the ecological strategy of each cytotype in relation to climate. All traits were measured in both years of the experiment unless otherwise noted (see Methods).

**Figure 3:**
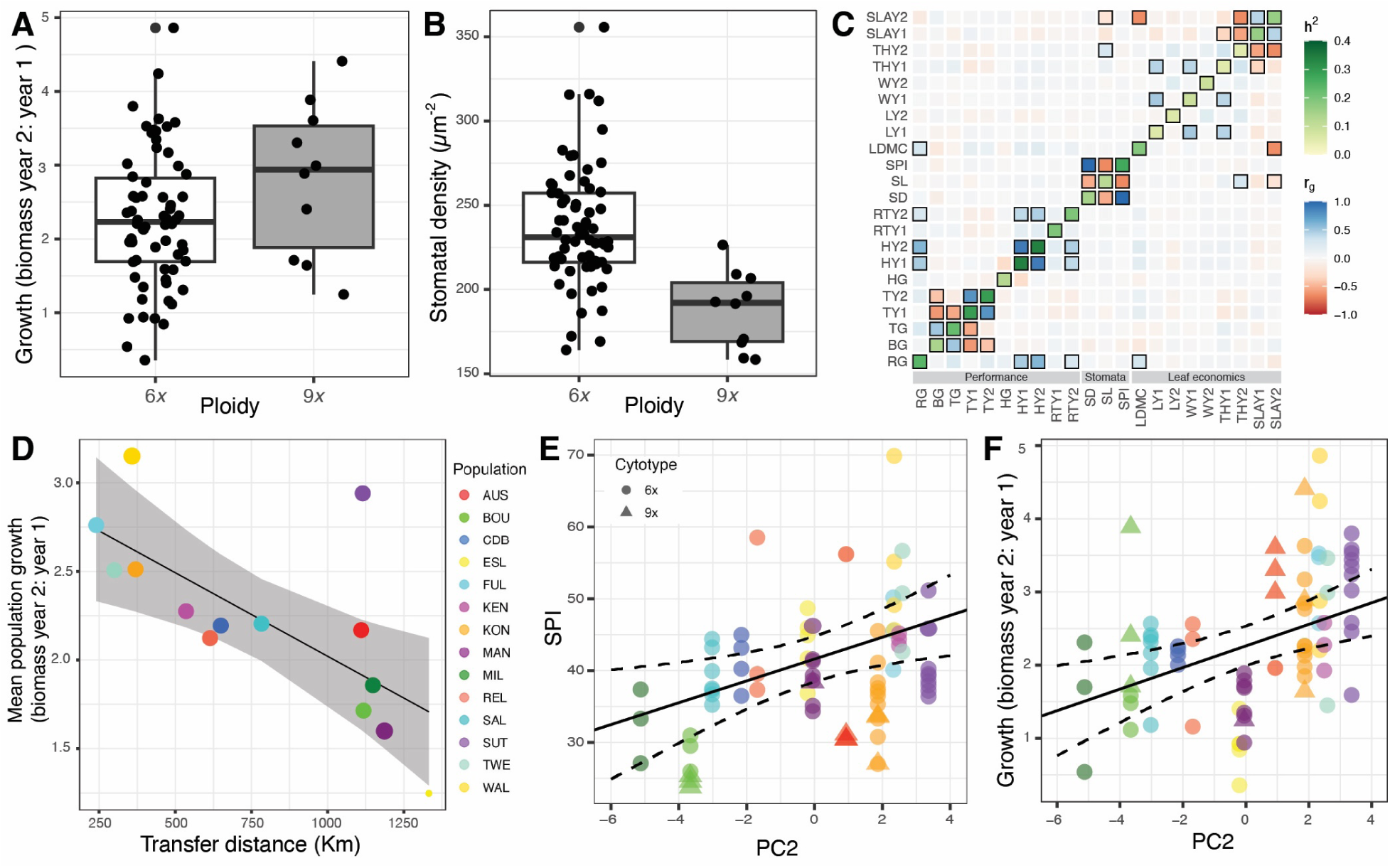
P**o**lyploidy **significantly effects growth and stomatal traits. (A)** The mean change in aboveground biomass and **(B)** stomatal density for each genotype by cytotype. **(C)** The estimated genetic correlation (*r_g_*) matrix between all measured phenotypes in 6*x* genotypes is plotted with the estimated narrow-sense heritability (*h*^2^) of each trait on the diagonal. Genetic correlations were estimated independently for each year where appropriate (year 1 is Y1 and year 2 is Y2). Axis labels refer to abbreviations for the measured phenotypes: change in aboveground biomass (RG), growth in basal area (BG), growth in number of tillers (TG), number of tillers (T), growth in height (HG), height (H, cm), percent of tillers flowering (RT), stomatal density (SD, *µ*m*^−^*^2^), stomata length (SL, *µ*m), stomatal pore index (SPI), leaf dry matter content (LDMC, mg/g), leaf length (L, cm), leaf width (W, cm), leaf thickness (TH, mm) and specific leaf area (SLA). Tiles with a black border have a 95% credible interval that does not cross zero. **(D)** The effect of geographic transfer distance on population mean change in aboveground biomass (i.e. growth). The gray area indicates the 95% confidence interval of the predicted values and the estimated mean RG of each population is overlaid as dots. **(E)** The predicted effect of PC2 on SPI (unitless) and **(F)** change in aboveground biomass from Model 2. The dashed lines are two standard errors from the mean and the average SPI is plotted for each genotype.

A PCA of best linear unbiased predictions (BLUPs) estimated for each genotype across all phenotypes did not identify clusters that would indicate the presence of ecotypes and did not support distinct morphologies for cytotypes or the East-West groups identified from SNPs (Fig. S10).

When phenotypes were individually examined, we found cytotype had a significant effect on between-year change in aboveground biomass (hereafter referred to as growth) and stomatal traits, but not leaf economic or other performance traits. The 9*x* cytotype grew approximately 45% more than the 6*x* cytotype (change in aboveground biomass; 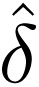= 1.02, 95% CI [0.422, 1.61], p = 0.0008) and had a lower stomatal pore index (SPI; 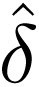= 8.63, 95% CI [4.28, 13], p = 0.0001) compared to the 6*x* cytotype (Fig. 3A). The lower SPI, which relates to capacity for gas exchange on the leaf surface (Sack *et al*. 2003), is consistent with the 9*x* cytotype exhibiting significantly larger stomata 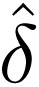=-4.3 *µ*m, 95% CI [−2.38, −4.40], p = *<* 0.0001) and lower stomatal density (40 *µ*m*^−^*^2^, 95% CI [19.4, 60.7], p = 0.0001; Fig. 3B). This pattern is consistent with a well-documented pattern of stomata size increasing with genome size (Beaulieu *et al*. 2008). Cytotype did not have a significant effect on any other individual phenotype (see SI Appendix for an extended discussion of the phenotype data).

The observed changes in both growth and stomatal traits may be correlated through increased DNA content leading to increased cell size (Doyle and Coate 2019; Clo and Kolář 2021) or may reflect trade-offs with other phenotypes. Growth and stomatal traits showed no significant genetic correlation (Fig. 3C), however, lending no evidence to a shared genetic mechanism outside of polyploidy. Rather, stomatal traits and growth exhibited trade-offs with other measured phenotypes (Fig. 3C). Stomatal density and stomata length were negatively genetically correlated (r*_g_* =-0.53) and showed opposing correlations with SPI (r*_g_* = 0.98, r*_g_* =-0.67). This is a common pattern in C_4_ grasses and indicates coordination of the ratio of stomatal pore area to leaf surface area, allowing grasses to optimize rates of photosynthesis and water loss (Taylor *et al*. 2012). Additionally, we found growth was positively genetically correlated with plant height in both years (*r_g,Y_*_1_ = 0.16, *r_g,Y_*_2_ = 0.19) and leaf dry matter content (only measured in year 2; *r_g_* = 0.28; Fig. 3C), which is a proxy of tissue density and mechanical resistance (Wright *et al*. 2004). The correlation between growth and leaf dry matter content suggests the observed change in biomass may be partially explained by investment in tissue density and support for greater plant height, as opposed to a greater number of tillers. Our results do not provide evidence for novel mechanisms that could explain these changes. Changes in DNA content and gene copy due to polyploidy, however, are well known to affect many of these traits (**?**Francis *et al*. 2008; Clo and Kolář 2021).

### Polyploidy may confer environmental adaptation to xeric climates

We hypothesized the observed differences in stomatal traits and growth rate among cytotypes may provide an advantage to the 9*x* cytotype in environments with low precipitation and high temperature variability, where the 9*x* cytotype is most abundant (McAllister *et al*. 2015). To test this hypothesis, we first used a PCA of 28 climate variables (Fig. S11) to characterize the primary axes of environmental variation across our sampled populations. The first two principal components described the majority of variation (65% and 26.3%; Fig. S11). Environmental PC1 was primarily driven by positive correlations with variables describing temperature and length of the growing season (mean annual temperature *r* = 99.1%, yearly growing degree days at 10◦C *r* = 95.4%), whereas PC2 was driven by positive correlations with precipitation variables (mean annual precipitation *r* = 91.3%).

We then tested whether or not the sampled *A. gerardi* populations exhibited patterns of environmental adaptation. To do so, we regressed population mean change in aboveground biomass (i.e. mean growth) against the geographic distance each population was transferred to the common garden (transfer distance) as a proxy for overall environmental similarity. If populations are adapted to environmental conditions similar to their home site, growth will decline as environmental similarity decreases. We found change in aboveground biomass steeply declines with increasing transfer distance (*r* =-0.69, p = 0.0064, Fig. 3D), suggesting *A. gerardi* populations perform best in environments similar to their population of origin. We also evaluated whether populations demonstrated adaptation to mean annual precipitation (MAP) and temperature (MAT), the primary axes of environmental variation identified in the environmental PCA. We use MAP and MAT instead of the PCs because they could be directly calculated for the common garden site in 2021 and 2022 while the PCs represent long-term climate averages. Similar to geographic distance, growth significantly declined with increasing difference in mean annual temperature; most sites were cooler than the common garden and individuals from colder sites grew more slowly (*r* =-0.61, p *<* 0.05; Fig. S13), indicating possible temperature stress. In contrast, we found that although there was a significant correlation between growth and the precipitation difference from the common garden (*r* =-0.82, p *<* 0.05; Fig. S13), the effect was asymmetrical — individuals from sites wetter than the common garden grew faster than individuals from drier sites. The asymmetry may result from populations in drier climates having evolved a conservative slow-growth strategy that cannot be hastened by simply increasing water availability (Reich 2014; Galliart *et al*. 2020). Overall, the populations with highest growth had home environments with temperatures similar to the common garden, but with greater water availability (Fig. S12).

After establishing that the sampled *A. gerardi* populations adapted to their home environments, we evaluated whether environmental variation is shaping phenotypic variation. We used linear mixed models to test the effect of environmental PC1 and PC2, while controlling for population structure and ploidy. As the traits were measured in a controlled environment, a significant effect of the home environment rejects a neutral model in which the environment does not affect phenotypic variation. We found that both PC1 and PC2 had a significant effect on numerous traits (see SI Appendix for an extended discussion of the phenotype data). Both environmental PCs had a significant effect on stomatal traits and growth, except that PC2 did not have a significant effect on stomatal length. Stomatal pore index (SPI), an index of stomatal pore area per leaf area calculated as the ratio of stomatal density to the square root of mean stomatal length, significantly increases with PC1 and PC2 (Fig. 3F, S31; PC1 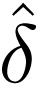= 0.05488, SE = 0.3394, p = 0.0011, PC2 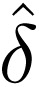= 1.50351, SE = 0.56940, p *<* 0.0001). This relationship is driven by a significant increase in stomatal density (Fig. S30; PC1 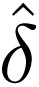= 0.6786, SE = 1.524, p = 0.0002, PC2 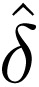= 6.960, SE = 2.505, p *<* 0.0001), and a somewhat weaker increase of stomata length (Fig. S29; PC1 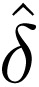= 0.1797, SE = 0.1658, p *<*0.0001). Together, this indicates populations in regions with higher mean annual temperatures and mean annual precipitation have greater stomatal pore area to leaf area. With increased relative stomatal pore area comes a trade-off of greater photosynthetic capacity but increased water loss, which may not be a risk if water is plentiful (Hetherington and Woodward 2003).

Consistent with greater photosynthetic capacity, we found growth significantly increases in populations from home environments with higher mean annual temperatures (Fig. S28; PC1 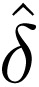= 0.0923, SE = 0.0276, p *<* 0.0001) and greater mean annual precipitation (Fig. 3F; PC2 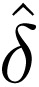= 0.144, SE = 0.0454, p *<* 0.0001), consistent with the transfer distance analysis described above. Notably, the stomatal and growth phenotypes of the 9*x* cytotype are most similar to populations from home environments with drier or shorter growing seasons, such as the most western and northern populations, suggesting polyploidy may provide an advantage in those populations. As 9*x* are most abundant in environments with reduced summer precipitation and high temperature seasonality (McAllister *et al*. 2015), this supports our hypothesis that the abundance of 9*x* is partially explained by polyploidy conferring adaptive phenotypes in some environments.

## Discussion

### Polyploidization in *A. gerardi* is common and recurrent

The high prevalence of polyploid species across North American grasslands suggests polyploidy may play an important role in the ecology and evolution of this ecosystem. Widespread polyploidy may result from demographic processes encouraging polyploid formation (Stebbins 1985) or evolutionary novelty generated during polyploidization (Van de Peer *et al*. 2017). Both mechanisms may interact with the environment to impact the probability of polyploidization and establishment. We tested these hypotheses by uniting genetic and phenotypic data across 29 populations of *A. gerardi*, the dominant species in endangered North American tallgrass prairies.

Leveraging a novel, haplotype-resolved, and chromosome-scale reference genome and the dynamics of transposable element insertion over time, we found the *A. gerardi* 6*x* cytotype formed through two poly-ploidization events occurring a minimum of 2.3 and 1.88 Mya (Fig. 1). This estimate is more recent than phylogenetic estimates of the divergence of hexaploid projenitors using four nuclear loci (8.6 Mya; Estep *et al*. 2014), and a recent preprint using similar methods (3.87 Mya; Stitzer *et al*. 2025). Our approach differed in the calculation of divergence (Ks), where Stitzer *et al*. (2025) estimated the mean *K_S_* within multi-species alignment blocks in order to compare divergence times across Andropogoneae species with less contiguous genome assemblies. Our analysis improved upon this approach by directly estimating the minimum time since both the tetra-and hexaploid formation from shared TE activity. The substantially more recent time we estimated for the hexaploid formation (1.88 Mya) suggests the hexaploidy may have arisen as recently as the early Pleistocene as the glacial cycles began. Our results are consistent with Stebbins (1985), who hypothesized the reduced habitat from glacial expansion forced the progenitor species of polyploids into closer proximity, increasing the probability of a successful secondary contact and allopolyploidization.

The ongoing polyploidization in *A. gerardi*, where neither the 6*x* or 9*x* cytotype has reached fixation, enabled us to examine the frequency of WGD in contemporary North American grasslands. We found the 9*x* cytotype is not genetically differentiated from the 6*x* cytotype (Fig. 2B-C, S8, S9), even within populations (Fig. 2E). The observed lack of differentiation is unlikely to be due to gene flow among 9*x* or between cytotypes due to irregular meiosis, unbalanced gametes, and low seed viability exhibited by the 9*x* cytotype (Norrmann *et al*. 1997; Tompkins *et al*. 2015; Norrmann and Keeler 2003). Indeed, the rare viable offspring of 9*x* genotypes that germinate in greenhouse settings are typically short-lived aneuploids and found in wild populations at less than 5% frequency range-wide (Norrmann *et al*. 1997; Keeler 1992; McAllister *et al*. 2015). Our results are consistent with each 9*x* genotype being the product of independent WGD events from the union of reduced and unreduced 6*x* gamete, and suggest many more origins of WGD than previous estimates (McAllister *et al*. 2015).

Other wild neo-autopolyploid systems have been documented but all have undergone multiple generations of sexual reproduction (Salony *et al*. 2024; Ramsey 2011). Neopolyploids, such as *A. gerardi*, are important resources for the continued study of the phenotypic, genetic, and ecological consequences of WGD in naturally formed polyploids (Edger *et al*. 2025).

Like the origin of the 6*x* cytotype, the recurrent formation of the 9*x* cytotype is a result of opportunity, yet it remains unclear whether WGD are constant or occur in environment-induced bursts. Unreduced gametes have been estimated to form at a rate of less than 1% in autopolyploid plants (Ramsey and Schemske 1998; Harlan and deWet 1975), but may be more common under stress (Wang *et al*. 2017; Sax 1936; Belling 1925). The severe’Dust Bowl’ drought, an extreme climatic event in the 1930’s (Schubert *et al*. 2004), may have resulted in such a burst. During this drought, there was extreme loss of North American C_4_ grasses, with up to a 70% loss of *A. gerardi* is some regions (Weaver and Albertson 1943), followed by a post-drought recolonization providing opportunity for polyploid establishment with reduced diploid competition.

### Neopolyploidy confers novel, adaptive phenotypic variation

Although increased opportunity for WGD has contributed to the abundance of the 9*x* cytotype, abundance of the 9*x* cytotype could be aided by novel phenotypic variation induced by WGD (Van de Peer *et al*. 2017). To test this hypothesis, we assessed whether the dominance of the 9*x* cytotype in xeric environments (McAllister *et al*. 2015) could be the result of adaptive phenotypic differences by examining ecologically relevant traits in a common garden experiment. Multivariate phenotypic diversity did not distinguish the two cytotypes, reflecting difficulties in identifying the cytotypes in the field, nor did it distinguish previously described ecotypes (Fig. S10, Galliart *et al*. 2019; Gray *et al*. 2014). Individual examination of phenotypes, however, revealed the cytotypes differ in important growth and stomatal traits (Fig. 3). The 9*x* cytotype had significantly greater change in aboveground biomass over two years, growing approximately 45% more than the 6*x*, and lower stomatal pore index (SPI).

To examine whether the WGD-induced variation may be beneficial, we tested if growth and stomatal traits were under selection by the environment using linear mixed models that controlled for neutral population structure. We found this to be true; both were significantly affected by temperature and growing season length (environmental PC1) and precipitation (environmental PC2) when controlling for ploidy and neutral population structure (Fig. 3E-F, SI Appendix). Growth and SPI increase with precipitation, temperature, and length of the growing season, consistent with previous *A. gerardi* common gardens along precipitation and photoperiod clines (McMillan 1959; Mendola *et al*. 2015; Galliart *et al*. 2020). Lower growth may reflect that a conservative growth strategy or overall smaller plant stature are beneficial in drier climates (Reich 2014). Low SPI has also been previously shown to be adaptive for *Andropogon* and C_4_ grasses in xeric climates as it is associated with lower maximum stomatal conductance and therefore decreased water loss (Taylor *et al*. 2012; Awada *et al*. 2002).

When considering the effect of polyploidy and the environment together, it becomes apparent that polyploidy may provide an advantage in xeric climates. We found WGD results in a reduction of SPI and therefore decreased water loss. Additionally, WGD increases growth which may provide a competitive advantage over the 6*x* cytotype in xeric-adapted populations when water is not limiting. As such, we propose that WGD induces phenotypes that contribute to 9*x* abundance in xeric climates. This may have trade-offs with population mean fitness due to low reproductive viability of 9*x* (Norrmann *et al*. 1997) despite having advantageous phenotypes. As a result, polyploidy may be a range-wide conservation concern in *A. gerardi* populations (Tompkins *et al*. 2015). Additional research into the cytotype-by-environment interaction is warranted to better understand the limits of where polyploids might establish. As *A. gerardi* is expected to decrease as much as 60% by 2070 without adaptation (Smith *et al*. 2017), polyploidy should be considered in the management and restoration of *A. gerardi* populations.

## Methods

### Sample collection

*A. gerardi* plants were either collected as rhizomes or grown from seed (Dataset 01). Necessary permissions and permits were obtained before collecting. Plants were sampled from 29 sites in the United States and Canada (Fig. 2A) and transported to the United States from Canada under phytosanitary certificate #3193417. Nearby sites were considered a single site in analyses resulting in a total of 25 sampled populations. Plants from rhizomes were collected following methods described in Phillips *et al*. (2023). Briefly, the plants were dug up with a shovel late in the growing season in 2016 through 2020. Any soil was washed off, the leaves were cut back to about 4 in in height to reduce transpiration, and the rhizomes were wrapped in wet paper towels for transportation to the Donald Danforth Plant Science Center in St. Louis, MO, USA. Plants grown from seed were grown from population bulk seed collected by Loretta Johnson at Kansas State University, KS, USA. The plants were potted and maintained in greenhouses with average conditions of 8°C day, 22°C night, 50% relative humidity, and a 16 hrs daylength. Once mature, the plants were maintained on outdoor benches year-round. The leaves were cut back and the pots were covered with straw mulch each winter. Voucher specimens were created for each population and have been deposited at the Missouri Botanical Garden (St. Louis, MO, USA.).

### Short-read sequencing of population panel

A set of 157 genotypes was processed for low-coverage sequencing at Cornell University (Dataset 01). Sequencing initially failed for 15 samples and was re-attemped at the University of California, Davis (UCD). DNA was extracted using approximately 100 mg of lyophilized leaf tissue and a DNeasy Plant Kit (Qiagen Inc., Germantown, MD). High throughput Illumina Nextera libraries were constructed and samples were sequenced with other plant samples in pools of 96 individuals in one lane of an S4 flowcell in an Illumina NovaSeq 6000 System with paired-end 150-bp reads, providing approximately 1.8x coverage for each sample from Cornell and 2.5x coverage from UCD.

A partially-overlapping additional subset of 50 genotypes were whole genome sequenced to 22-56x (mean = 37x) coverage by the Department of Energy (DOE) Joint Genome Institute (JGI) in Berkeley, California (Dataset 01). High molecular weight DNA was extracted from young tissue using the protocol of Doyle and Doyle (1987) with minor modifications. Flash-frozen young leaves were ground to a fine powder in a frozen mortar with liquid nitrogen followed by extraction in 2% CTAB buffer (that included proteinase K, PVP-40 and beta-mercaptoethanol) for 30 min to 1 h at 50°C. After centrifugation, the supernatant was extracted twice with 24:1 Chloroform:Isoamyl alcohol. The upper phase was transferred to a new tube and 1/10th volume 3 M Sodium acetate was added, mixed, and DNA precipitated with iso-propanol. DNA precipitate was collected by centrifugation, washed with 70% ethanol, air dried for 5-10 min and dissolved in an elution buffer at room temperature followed by RNAse treatment. DNA purity was measured with NanoDrop, DNA concentration measured with Qubit^TM^ HS kit (Invitrogen^TM^), and DNA size was validated by Femto Pulse System (Agilent). Sequencing libraries were constructed using an Illumina TruSeq DNA PCR-free library kit using standard protocols. Libraries were sequenced as above. The quality and possible contamination of the short-read sequences was evaluated to confirm taxonomy of the sequences (see SI Appendix). 9 samples (6%) were identified as anything other than *Schizachyrium* or *Andropogon* and were discarded.

### Genome size estimation

We estimated genome size with flow cytometry for all sampled individuals, except for a a few plants that died before estimation. Flow cytometry methods were previously described in Phillips *et al*. (2023). Briefly, maize B73 inbred line (5.16 pg*/*2C) was used as an internal standard, and we analyzed three replicate samples for each individual. The cell count, coefficient of variation of FL2-A, and mean FL2-A were recorded for the target and reference sample with no gating, and replicates were averaged to calculate genome size (Dataset 02).

Of the individuals for which genome size couldn’t be estimated with flow cytometry, sixteen had sufficient sequencing coverage for ploidy to be determined using nQuire (Weiß *et al*. 2018). Thirty high-coverage genotypes with genome sizes successfully estimated using flow cytometry were included in the nQuire analysis as controls. Of the 30 control genotypes, 29 were 6*x* and one was 9*x*. nQuire uses a Gaussian Mixture Model to model the read frequency histogram expected for diploids, triploids, and tetraploids. As all three subgenomes are resolved in the *A. gerardi* genome described below, we expect the 6*x* genotypes to have a diploid distribution and the 9*x* genotypes to have a triploid distribution. Maximized log-likelihoods were estimated for each genotype and normalized to the maximized log-likelihood of the data modeled under a free model. The ploidy model with the highest normalized log-likelihood was assigned the ploidy of the genotype. To assess noise and error in this method, the normalized maximized log-likelihoods for each of the three ploidy models were plotted against each other in R (v4.2.2, R Core Team 2017; Figure S14).

Of the remaining individuals for which genome size could not be estimated with flow cytometry or sequenced-based approaches, ploidy could be inferred for one population based on a previous study. The population CUI (Cuivre River State Park, MO, USA) was previously sampled for cytotypic composition by McAllister *et al*. (2015) and found to be 100% 6*x*. Given this study is relatively recent, we assumed all individuals sampled from CUI are 6*x*.

### Genome sequencing for the *A. gerardi* reference genome

We sequenced *A. gerardi* (collection number *Kellogg-1272*) using a whole genome shotgun sequencing strategy and standard sequencing protocols. Sequencing reads were collected using Illumina and PacBio platforms. Illumina and PacBio reads were sequenced at DOE JGI and the HudsonAlpha Institute in Huntsville, Alabama. Illumina reads were sequenced using the Illumina NovoSeq 6000 platform, and the PacBio reads were sequenced using the SEQUEL II platform. One 400 bp insert 2×250 Illumina fragment library (92.80x coverage) was sequenced along with one 2×150 OmniC library (54.38x; Table S1). Prior to assembly, Illumina fragment reads were screened for PhiX contamination. Reads composed of *>*95% simple sequences were removed. Illumina reads *<*50 bp after trimming for adapter and quality (q*<*20) were removed. The final read set consists of 1,601,302,817 reads for a total of 92.80x of high-quality Illumina bases. For the PacBio sequencing, the total circular consensus sequencing (CCS) sequence yield consisted of 16,785,606 reads (average size 19,680 bp) that produced 343.73 Gbp (85.93x; Table S2).

### Genome assembly and construction of pseudomolecule chromosome

All PacBio CCS reads were assembled using HiFiAsm+HIC assembler (v15.1, Cheng *et al*. 2021) and subsequently polished using the 1,601,302,817 Illumina fragment 2×250 reads (92.80x) to resolve homozygous SNP/indel errors with RANCON (v1.4.10; Vaser *et al*. 2017, Table S3-S6). This produced initial assemblies of both haplotypes. The initial haplotype 1 (HAP1) assembly consisted of 1,064 scaffolds, with a contig N50 of 55.6 Mbp, and a total genome size of 2,780.5 Mbp (Table S3). The initial haplotype 2 (HAP2) assembly consisted of 731 scaffolds, with a contig N50 of 59.9 Mbp, and a total genome size of 2,701.5 Mbp (Table S4).

Hi-C Illumina reads (54.38x) from *A. gerardi* (collection number *Kellogg 1272*) were separately aligned to the HAP1 and HAP2 contig sets with Juicer (v1.8.9, Durand *et al*. 2016), and chromosome-scale scaffolding was performed with 3D-DNA (v180922, Dudchenko *et al*. 2017). No misjoins were identified in either the HAP1 or HAP2 assemblies. The contigs were then oriented, ordered, and joined together into 30 chromosomes per haplotype using the Hi-C data. A total of 38 joins were applied to the HAP1 assembly, and 38 joins for the HAP2 assembly. Each chromosome join is padded with 10,000 Ns. Contigs terminating in significant telomeric sequence were identified using the (TTTAGGG)n repeat, and care was taken to make sure that they were properly oriented in the production assembly.

Scaffolds that were not anchored in a chromosome were classified into bins depending on sequence content (SI Appendix). Contamination was identified using blastn against the NCBI non-redundant nucleotide collection (NR/NT) and blastx using a set of known microbial proteins. SNPs/indels were corrected in both haplotypes with 62x Illumina reads (Appendix SI).

The final version 1.0 HAP1 release contained 2,669.3 Mbp of sequence, consisting of 88 contigs with a contig N50 of 63.1 Mbp and a total of 99.90% of assembled bases in chromosomes. The final version 1.0 HAP2 release contained 2,588.3 Mbp of sequence, consisting of 68 contigs with a contig N50 of 59.2 Mbp and a total of 99.93% of assembled bases in chromosomes.

Completeness of the gene space of both haplotypes was measured with Benchmarking Universal Single-Copy Orthologs (BUSCO; Manni *et al*. (2021)) using the Embryophyta and Eukaryota (OrthoDB v9) profiles.

### Subgenome identification and divergence

Following Session and Rokhsar (2023), the three subgenomes were then clustered into groups of 10 chromosomes based on shared enriched 12-mer content. For each triplet of chromosomes, all 12-mers were identified from a frequency of 20 – 5000, with a minimum of 50 occurrences being required on one of the triplets. A positively enriched 12-mer had one of the three with at least 3 times the count of the other two. All occurrences of the enriched 12-mers were counted across the chromosomes and converted to a binary call.

Binary k-mer calls were subset to sites observed in ≥5 chromosomes and used to construct a symmetric binary distance matrix. Clustering was then accomplished on the distance matrix by partitioning around medoids (PAM) with the R package cluster (v2.1.4, Maechler *et al*. 2012).

The 3 subgenomes were designated as A, B, and C. The”A” subgenome was identified as being most closely related to *A. virginicum* using a HipMer (Georganas *et al*. 2015) assembly of *A. virginicum* to mask the 3 subgenomes using 18-mers. The A-genome consistently shared more content with *A. virginicum* than the other two subgenomes. Chromosomes were numbered and oriented within the 3 subgenomes using *Sorghum bicolor*, and the resulting sequence was screened for retained vectors and contaminants (Tables S5, S6).

To estimate divergence among subgenomes, we identify subgenome-specific 17-mers found in at least 40 copies with SubPhaser (-k 17-q 40-baseline-1), as well as subgenome-specific LTR retrotransposons (Jia *et al*. 2022). To convert substitutions per site to a mutation rate, we used the grass-specific rate of 6.5 × 10*^−^*^9^ (Gaut *et al*. 1996). While this does not completely account for differences in generation time among perennial lineages, it is more appropriate than comparable estimates derived from annual grasses like maize or rice.

We generated gene trees of syntenic loci within *A. gerardi*. First, we defined syntenic positions using Anchorwave (Song *et al*. 2022), using *Paspalum vaginatum* (Sun *et al*. 2022) genes as reference and each *A. gerardi* haplotype assembly as a query. We set-R 3-Q 1 to allow for the *A. gerardi* hexaploidy relative to diploid *P. vaginatum*, such that each *P. vaginatum* gene can participate in up to three syntenic blocks in *A. gerardi*. We extracted loci found in all six haplotypes, aligned these together with *P. vaginatum* sequence using mafft (--genafpair --maxiterate 1000 --adjustdirection; Katoh and Standley 2013), and built trees of each locus using RAxML (-m GTRGAMMA-p 12345-x 12345-# 100-f a; Stamatakis 2014. We summarized these gene trees using ape (Paradis and Schliep 2019) in R, rooting with *P. vaginatum*, then extracting the underlying topology relating each haplotypic and subgenomic copy (e.g. (((A1,A2),(B1,B2)),(C1,C2));). We also measured pairwise synonymous divergence (*K_S_*) for all six *A. gerardi* copies at each locus using seqinR (Charif and Lobry 2007).

### Genome annotation

Transcript assemblies were made for each haplotype from about 1.2 billion pairs of 2×150 stranded paired-end Illumina RNA-seq reads using PERTRAN, which conducts genome-guided transcriptome short read assembly via GSNAP (Wu and Nacu 2010) and builds splice alignment graphs after alignment validation, realignment, and correction. Approximately 14.8 million PacBio Iso-Seq CCS reads were corrected and collapsed by a genome-guided correction pipeline, which aligns CCS reads to the respective haplotype with GMAP (Wu and Nacu 2010) and corrects introns for small indels in splice junctions when all introns are the same or 95% overlap for single exon, to obtain about 774,000 and 766,000 putative full-length transcripts for HAP1 and HAP2, respectively.

Subsequently, 829,257 (HAP1) and 820,009 (HAP2) transcript assemblies were constructed using PASA (Haas *et al*. 2003) from the RNA-seq transcript assemblies and the respective haplotype. The PASA-improved gene model proteins were subject to protein homology analysis to the above-mentioned proteomes to obtain the C-score, a protein BLASTP score ratio to the mutual best hit BLASTP score, and protein coverage, the highest percentage of protein aligned to the best of homologs, for each transcript. PASA-improved transcripts were selected if their C-score was ≥0.5 and protein coverage ≥0.5, or it had EST coverage but its CDS overlapping with repeats is *<*20%. Gene models with CDS that overlap repeats more than 20% must have a C-score ≥0.9 and homology coverage ≥70% to be selected. Additionally, gene models were subject to Pfam analysis (Mistry *et al*. 2021) and gene models with *>*30% TE domains were removed. Gene models that were incomplete, had low homology support, a short single exon (*<*300 bp CDS) without a protein domain, nor good expression were manually filtered out. Transposable elements were annotated using the Extensive de novo TE Annotator (EDTA; Ou *et al*. 2019).

### Variant calling and genotyping

The quality of the raw WGS short read sequence data was assessed using FastQC (v0.11.6, http://www.bioinformatics.babraham.ac.uk/projects/fastqc/). Samples re-sequenced at UCD were trimmed using fastp to remove adapters, polyG tails, and the first 9 bp on each read while requiring reads to have a minimum length of 36 bp (fastp-l 36-Q --trim front1 9 --trim front2 9; v0.20.1, Chen *et al*. 2018). Samples sequenced at JGI and Cornell did not require trimming to improve alignment quality.

The sequence data were aligned to the *A. gerardi* reference genome with bwa-mem2 (v2.2; Vasimuddin *et al*. 2019). Reads from samples that were sequenced on multiple lanes were combined into a single fasta file prior to alignment. The BAM files were sorted using SAMtools (v1.7; Danecek *et al*. 2021), read groups were added using Picard AddOrReplaceReadGroups, and duplicates were removed with Picard MarkDuplicates (v2.27, http://broadinstitute.github.io/picard) using default settings. Alignment quality was assessed using QualiMap (qualimap bamqc-nt 1000-nt 12-nw 400 --skip-duplicated; v.2.1.1, Okonechnikov *et al*. 2016). BAMs from genotypes that were independently sequenced multiple times were merged into a single BAM. High coverage samples sequenced by JGI were subsampled to approximately 2-4x coverage (samtools view-b-s 0.06). Genotypes with less than 90% reads mapping, *<*0.5x coverage, or missing genome size data were excluded from downstream analyses.

For analyses that included both 6*x* and 9*x* genotypes, we identified and filtered variable sites using BCFtools (Li 2011). Variable sites were called using BCFtools mpileup and BCFtools call (v1.16; Li 2011). Sites were filtered to exclude those that are multiallelic and have low mapping quality and sequencing quality with GATK (gatk VariantFiltration-filter “QUAL ≤ 30”-filter “MQ ≤ 30” and gatk SelectVariants--restrict-alleles-to BIALLELIC; v4.2; Van der Auwera and O’Connor 2020). Sites were additionally filtered for *<*20% missing data and a genotype depth ≥1 using a custom R script. A maximum depth filter was applied in order to exclude sites where paralogs may be mapping (Phillips 2024). We defined the maximum depth cutoff at each site as the 99th percentile, assuming coverage follows a Poisson distribution.

To address biases due to ploidy differences, we called single-read genotypes with ANGSD (Korneliussen *et al*. 2014; Phillips *et al*. 2023). Single-read genotypes were sampled at the filtered sites directly from the BAMs using ANGSD (angsd-doIBS 1-doMajorMinor 3-doCounts 1; v0.934; Korneliussen *et al*. 2014). Single-read genotypes were generated by randomly drawing a read at each site. If the read draw has the reference allele, the genotype is”1” while the alternate allele is”0”.

### Assessment of population structure and diversity

To assess population structure, we first randomly sampled one-hundred thousand sites to minimize the impacts of linkage disequillibrium. A principal component analysis (PCA) was run using ANGSD (angsddoCov 1). We additionally ran a PCA with PCangsd (Meisner and Albrechtsen 2018) on only the high-coverage 6*x* genotypes to ensure we were accurately capturing the broader population structure and diversity. To do so, genotype likelihoods were called with ANGSD (-GL 1) and the quality metrics described above were applied for filtering. We subset to 30,000 sites, to minimize the impacts of linkage disequilibrium, and ran PCangsd with default settings.

Population admixture was assessed by estimating the individual ancestry coefficients and number of genetic clusters (K) using the STRUCTURE admixture model (v2.3.4, Pritchard *et al*. 2000). STRUCTURE was run for a K of 2 through 24 for 3 replicates of 85,000 iterations per model (including a 10,000 burn-in).

We specified PLOIDY as 1 because the single-read genotypes only sample one haplotype. Convergence was confirmed by consistent results between replicates (Fig. S3).

Population differentiation was estimated with pairwise *F_ST_* and *ρ* (Ronfort *et al*. 1998; Meirmans *et al*. 2018). Values were estimated between the East and West genetic groups identified in the above analyses and pairwise between all cytotype-population combinations. For the pairwise population analysis, three genotypes were randomly selected from cytotype-population groups containing at least three genotypes. Then, population allele frequencies were calculated from the single-read genotypes. Subsequently, pairwise *F_ST_* was calculated as 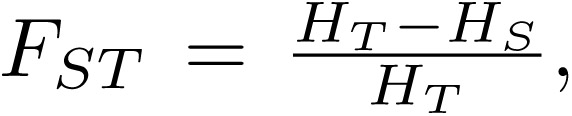 where *H_T_* is the expected heterozygosity and *H_S_* is the observed heterozygosity within the two populations. Pairwise *ρ* was calculated as 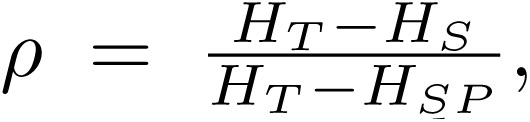 where *H_SP_*is the ploidy-corrected *H_S_*, following Meirmans *et al*. (2018). *F_ST_* and *ρ* were estimated per-site for each pairwise comparison and then averaged. To determine if within 9*x*, within 6*x*, and between cytotype genetic differentiation was significantly different, we estimated the mean *F_ST_* and *ρ* for the three comparison types (6*x*-6*x*, 9*x*-9*x*, 6*x*-9*x*) using the R function emmeans and tested whether the means differed using a Tukey pairwise comparison implemented in contrast (v1.1, Searle *et al*. 1980) at a 95% confidence level.

A kinship matrix was estimated following VanRaden (2008), which was found to be the best estimator for polyploid populations (Amadeu *et al*. 2020; Bilton *et al*. 2024), using a custom script in R. The diagonal elements were set to 1 for use in downstream analyses, as an accurate estimate of kinship within an individual cannot be made using a draw of a single read. Downstream analyses produced the same results given a diagonal of 1, 0, or random values. Clones were identified as having a kinship coefficient greater than 0.4; only two hexaploid genotypes were identified as clones. Population structure analyses were run with and without clones and results were similar.

### Estimation of hexaploid genetic diversity

Nucleotide diversity (*θ_P_*), Watterson’s theta (*θ_W_*), and Tajima’s D were estimated for all 6*x* genotypes in 10,000 and 50,000 bp windows using ANGSD (v0.935, Korneliussen *et al*. 2013). The folded site frequency spectrum (SFS), thetas, and Tajima’s D were estimated independently for each population. The two identified clones were excluded from estimation of hexaploid genetic diversity. Individual inbreeding coefficients were estimated only in 6*x* genotypes with high coverage WGS data, as low coverage data can result in significant bias (Bilton *et al*. 2024). The inbreeding coefficients were estimated in parallel for each chromo-some using ngsF (v1.2.0, Vieira *et al*. 2013). Results for all analyses were plotted in R using ggplot2 (v3.4; Wickham 2016).

### Common garden experimental design

A subset of 85 genotypes from 14 populations was planted in a common garden in Columbia, Missouri at the University of Missouri Genetics Farm in May 2021. The populations and genotypes were selected to maximize diversity in home environments and representation of the 9*x* cytotype. Additionally, the selected genotypes were required to have WGS and flow cytometry data. The selected genotypes were vegetatively propagated by splitting the rhizomes to produce three clonal replicates. Clonal replicates were bulked at the Donald Danforth Plant Science Center and UCD greenhouse facilities. Genotypes were planted in a randomized block design with three blocks, placing one random clonal replicate per genotype in each block (Fig. **??**). Random positions within the block were modified only when two genotypes from the same population were neighbors.

Prior to planting, the field was covered with landscaping cloth (DeWitt Sunbelt Woven Ground Cover) to prevent competition from annual grasses and broadleaf weeds. Plants were spaced 4.5 ft (1.37 m) apart in a grid pattern and the landscape cloth extended for at least 4.5 ft (1.37 m) around the edge of the blocks. The three blocks were separated by 9 ft (2.74 m). The holes cut in the landscaping cloth were cut in an X-shape at least 3 ft wide in each direction to ensure lateral growth was not constrained. After planting, the field was irrigated once to promote establishment and was not irrigated for the rest of the experiment, relying only on rainfall. The field was hand-weeded and broadleaf weeds were sprayed with 2,4-D herbicide early in the spring when the *A. gerardi* plants were small and likely to be outcompeted by weeds.

### Phenotyping leaf functional traits

The field was phenotyped in early September of 2021 and 2022. Six leaves were collected from each plant each year, selecting the youngest fully expanded leaves on 6 different tillers (Perez-Harguindeguy *et al*. 2016; Garnier *et al*. 2001). Leaves were cut at the ligule with scissors, placed in a plastic bag with a damp paper towel, then stored overnight in a 4*^◦^*C fridge before phenotyping. Within 72 hours of collection, fresh weight and leaf lamina thickness were measured. Fresh weight was only measured in 2022. Leaf lamina thickness was measured at the base of the leaf blade using digital calipers, taking care to avoid the midrib. The leaf blades were scanned for measurement of width, length, and one-sided area using ImageJ (v1.53-1.54; Schneider *et al*. 2012). Leaf length was measured as the length of the leaf b from the ligule to the leaf tip and leaf width was measured at the widest part of the leaf. Specific leaf area (SLA), leaf dry matter content (measured in 2022 only), and leaf density were calculated for each leaf (Vile *et al*. 2005; Garnier *et al*. 2001). After scanning, abaxial leaf impressions were taken on the fresh leaves using dental putty (Zhermack elite HT+ light body fast set) for analysis of stomatal traits. Three impressions were taken per leaf (bottom, middle, top) to capture developmental variation. After the dental putty impressions were taken, the fresh leaves were placed in a 7 in manilla envelope and dried at 70*^◦^*C for at least 72 hr. After the leaves were dried, dry mass was measured.

We measured stomatal traits by taking a negative of the dental putty impression with clear nail polish, which was imaged at 10X magnification using a Leica DM1000 microscope and MC170 HD digital camera. Using ImageJ, guard cell length was measured along the longest portion of the guard cell for five stomata per impression. Additionally, stomatal density was measured for each impression. As stomatal density is very high in *A. gerardi*, the image was cropped to a smaller area containing at least 10 stomata. In the cropped image, the area and number of stomata were measured. SPI was measured as stomatal density divided by the square root of mean guard cell length. Stomatal traits were measured in both years but data were incomplete in 2021. As a result, we discuss only the 2022 data. We tested for differences in mean stomatal traits between each group of impressions along the leaf using a Tukey test with a 95% confidence interval implemented in emmeans and contrast (v1.1, Searle *et al*. 1980). Results were similar across leaf impressions (Fig. S15).

### Phenotyping performance traits

Survival was recorded each year, although overall mortality was low (11%). Additionally, we measured the number of tillers, percent of tillers flowering, plant height, basal area, and aboveground biomass. Plant height was measured as the length of the longest tiller from the ground to the tip of the inflorescence. To estimate basal area, we measured the diameter of the base in two perpendicular directions and then calculated the area as an ellipse (Aspinwall *et al*. 2013). After all phenotypes were collected for the year, aboveground biomass was cut off approximately 4 in above the crown using grass shears. The biomass was placed in a brown paper bag and dried at 37*^◦^*C for 3-5 days until the dry weight stabilized. Once dry, the total dry aboveground biomass was measured. Root and below-ground traits were not measured as they required destructive sampling. Basal growth, tiller growth, and relative growth were calculated as the ratio between year 2 and year 1 measurements for basal area, number of tillers, and aboveground biomass, respectively.

### Trait data analysis

The effect of ploidy and the environment on trait variation was tested using two linear mixed models built in the R package Sommer (v4.3.3, Giovanny 2016). First, the effect of ploidy on a given phenotype (*Y*) was tested with Model 1, where ploidy (*P*) was specified as a fixed effect and population (*z*) and genotype (*g*) were random effects. Covariance among genotypes was specified by the kinship matrix (**K**). Year (*t*) was included as a random effect if the phenotype was measured in multiple years.

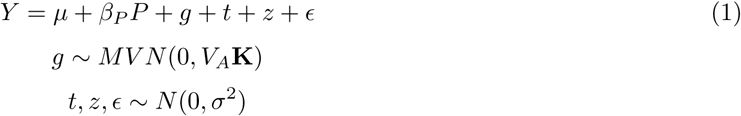

To test the effect of a population’s home environment on trait variation, yearly climate data for each population and the common garden site was extracted from ClimateNA using ClimateNAr (v1.1, Wang *et al*. 2016). The yearly ClimateNA variables (Fig. S11B) were averaged from 1961 to 1990. The average growing degree days per year with a base temperature of 10*^◦^*C (GDD) were separately estimated using daymetr (v1.7, Hufkens *et al*. 2018) for the period of 1980 to 2000. A PCA was run on the climate data to describe the environmental distance between populations using prcomp(center = T, scale = T) in R. PC1 and PC2 were selected to describe the home environment as they described the majority of environmental variation (Fig. S11). PC1 (*E*_1_) and PC2 (*E*_2_) were specified as fixed effects and added to the previously described model:

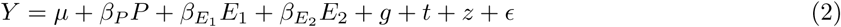

The residuals of each model were qualitatively assessed for normality, homogeneity of variances, and independence. Transformations were applied where needed; a log, exponential, and inverse normal transformation were applied to basal growth, percent of tillers flowering, and SLA, respectively. The significance of the fixed effects were tested with an ANOVA. Significance was evaluated for all tests at a confidence (alpha) of 95%. A Bonferroni multiple test correction was applied and effects were considered significant when the p-value was less than 0.0033.

The genetic correlation and heritability of the phenotypes were estimated with MegaLMM (v0.1.0,Runcie *et al*. 2021) using Model 1 without year in the model. Rather, we treated traits measured in multiple years separately. We specified 20 latent factors and used the default priors. We extracted the posterior means for lambda, genetic variance, and genetic covariances from a single model run for 1000 iterations after a burn-in of 500 iterations. We also estimated the 95% credible interval of the posterior distributions for the genetic covariances and heritabilities.

Change in aboveground biomass was regressed against geographic and climate transfer distance to assess location adaptation. Using Sommer, best linear unbiased predictors (BLUPs) were estimated for each population using Model 1. The grand mean was added to the estimated BLUPs to aid interpretation. Geographic transfer distance was estimated as the Haversine distance between the population and common garden using geodist (v0.0.8, Padgham 2021). Climate distance was measured by the difference in average mean annual precipitation (MAP) and mean annual temperature (MAT) between the common garden environment and home environment. Average MAP and MAT was estimated for each population from the ClimateNA data described above. Daily precipitation and temperature data for the common garden in 2021 and 2022 was downloaded for the Columbia-Jefferson Farm and Gardens (Boone County, MO, USA) weather station from the Missouri Historical Agricultural Weather Database. The average daily temperature was averaged across years to estimate the common garden MAT. Total daily precipitation was summed for each year, then averaged to estimate the common garden MAP. Finally, the three measures of transfer distance (geographic, MAP, and MAT) were regressed against population BLUPS with lm in R.

## Data availability

The sequencing data is available on NCBI Sequence Read Archive (SRA) under BioProject PRJNA1109389 and PRJNA519007. The genotype metadata, raw phenotype data, and flow cytometry data are available on Dryad at https://doi.org/10.5061/dryad.gxd2547v1. The *A. gerardi* reference genome is available on Phytozome under genome IDs 784 and 783. All scripts for genotype calling, population genetic analyses, and trait data analysis can be found at https://github.com/phillipsar2/andro_snakemake.

## Acknowledgements

This project was funded by the National Science Foundation (NSF) grant numbers 1822330 and 1934384, the Davis Botanical Society, and the Botanical Society of America. JRI also acknowledges funding from USDA Hatch project CA-D-PLS-2066-H 548. The work (proposal: 10.46936/10.25585/60001091) conducted by the U.S. Department of Energy Joint Genome Institute (https://ror.org/04xm1d337), a DOE Office of Science User Facility, is supported by the Office of Science of the U.S. Department of Energy operated under Contract No. DE-AC02-05CH11231. JTL is supported by the Center for Bioenergy Innovation (CBI), U.S. Department of Energy, Office of Science, Biological and Environmental Research Program under Award Number ERKP886. This research used the High-Performance Computing Core Facility (HPC@UCD) and DNA Technologies and Expression Analysis Core at the University of California, Davis, and the Genomics Facility (RRID: SCR 021727) of the Biotechnology Resource Center of Cornell Institute of Biotechnology. We would like to thank Christine McAllister, Michael McKain, and Loretta Johnson for providing germplasm and Jacob Washburn, Kate Guill, Norman Best, Jim Elder, Susan Melia-Hancock, Nancy Salazar-Vidal, Abiskar Gyawali, Melanie Carraher, Grace Sidberry, and Christopher Browne for assistance in planting and maintaining the common garden. We would also like to thank the agencies and people that provided permits and aided collections: Matt McCaw and the City of Austin Water Quality Protection Lands (WQPL), North Carolina Department of Agriculture and Consumer Services, NC Plant Conservation Program, the Suther family and Rev. Dennis Testerman, Chris Matson and Florida Park Service and Department of Environmental Protection, Kyle Dillard and the Milnesand Prairie Preserve (Creamer Ranch), City of Boulder Open Space and Mountain Parks, Lynn Riedel, Brian Anacker, Bess Bookout, Tim Teetaert and the Manitoba Tallgrass Prairie Preserve, Robert D. Bradley, Ken McCarty and Cuiver River State Park, Malissa Briggler and Victoria Glades Conservation Area. Finally, we would like to acknowledge Felix Andrews for helpful discussion about the relative semantic merits of nonoploid, triploid, and enneaploid terminology.

