## Supplementary material for "The origins and adaptive consequences of polyploidy in a dominant prairie grass": SI Appendix

**This PDF file includes:**

- Figs. S1 to S16
- Tables S1 to S6
- Supplementary Information Text
- SI References

### Supplementary Figures

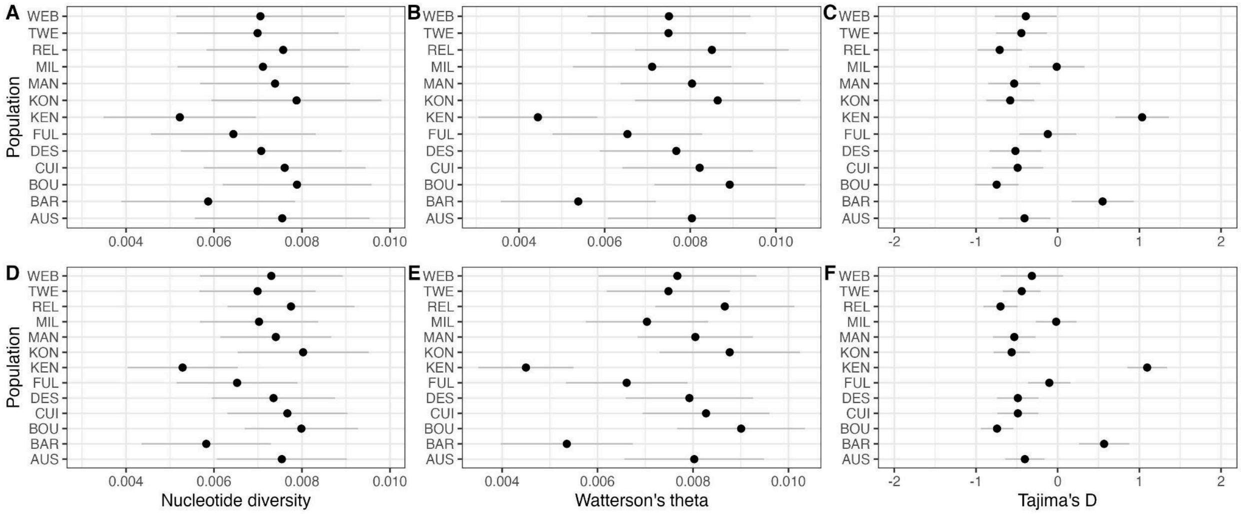


**Figure S1: Population estimates of hexaploid genetic diversity.** Nucleotide diversity (A, D), Watterson's theta (B, E), and Tajima's D (C, F) were estimated in 10 Kbp (A, B, C) and 50 Kbp (D, E, F) windows. The mean values for each population are shown as a black dot with an error bar indicating one standard deviation from the mean.

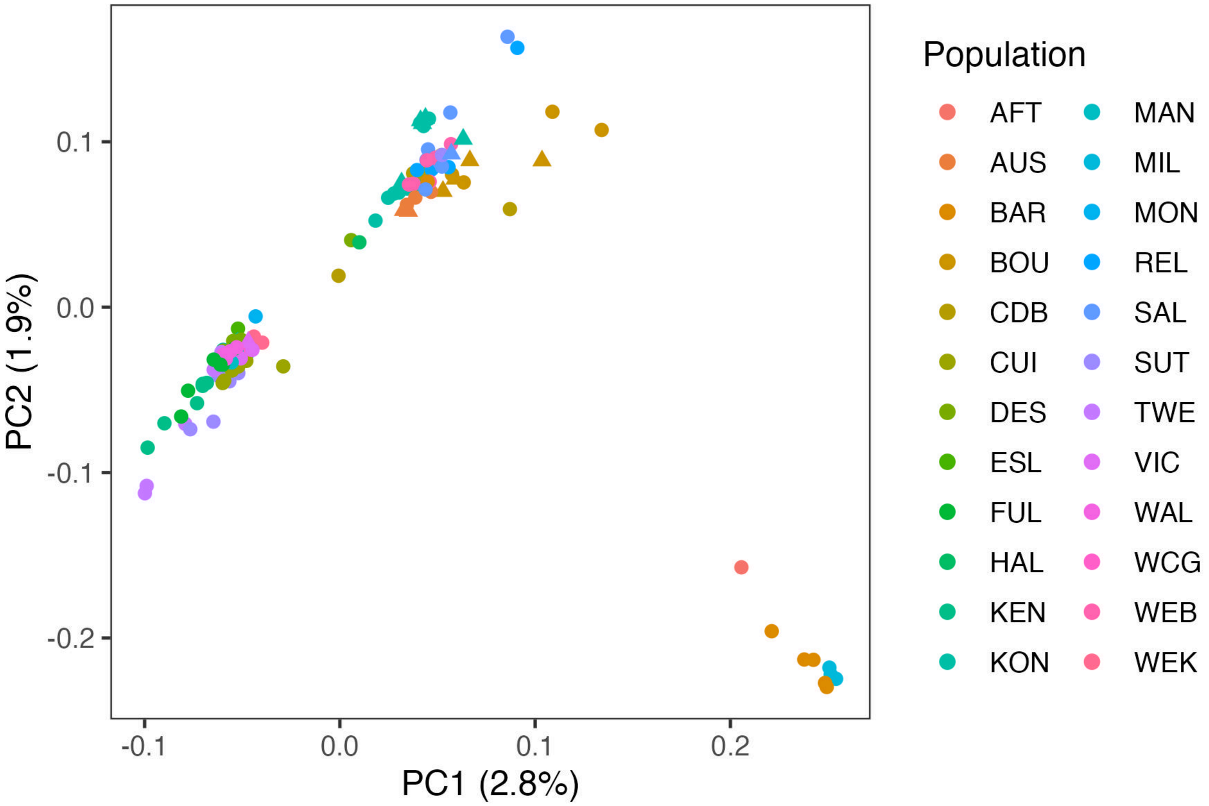


**Figure S2: Principal component analysis of single read genotypes for all sequenced genotypes.** The first two principal components are plotted for each genotype with the color of each point indicating the population of origin. The shape of each point indicates the ploidy of the sample where 6*x* are circles and 9*x* are triangles**.**

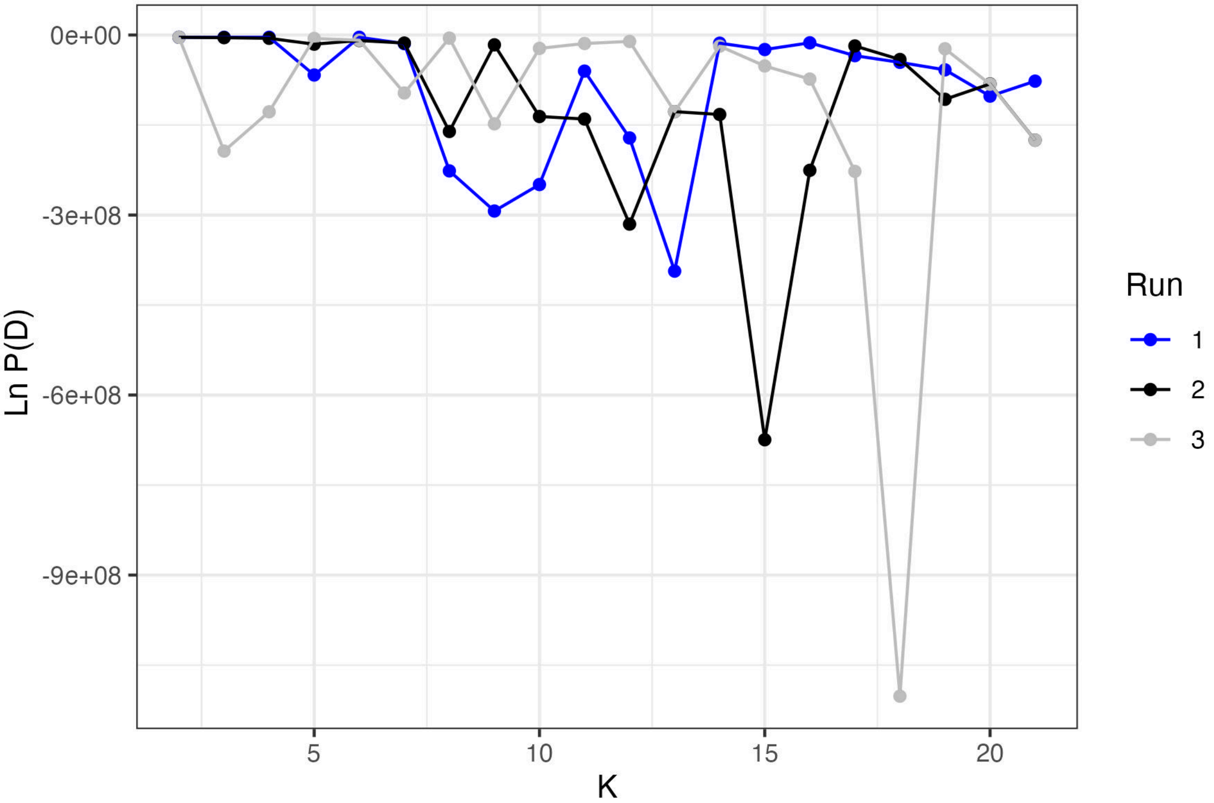


**Figure S3: The estimated log probability of the sequence data given K admixture groups as estimated by STRUCTURE.** Three runs were estimated for each K. The estimated probability and consistency across runs declines with an increasing value of K.

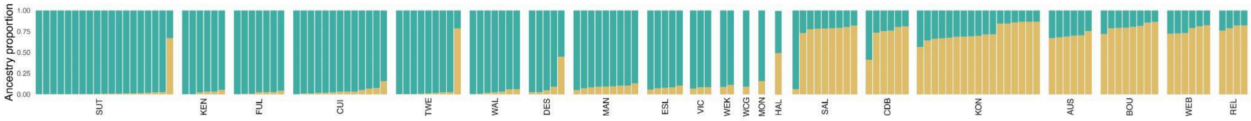


**Figure S4: STRUCTURE results when run without the BAR, MIL, and AUS populations.** Two runs were completed for each K, and had similar results. Colors and population ID follow Fig. 2.

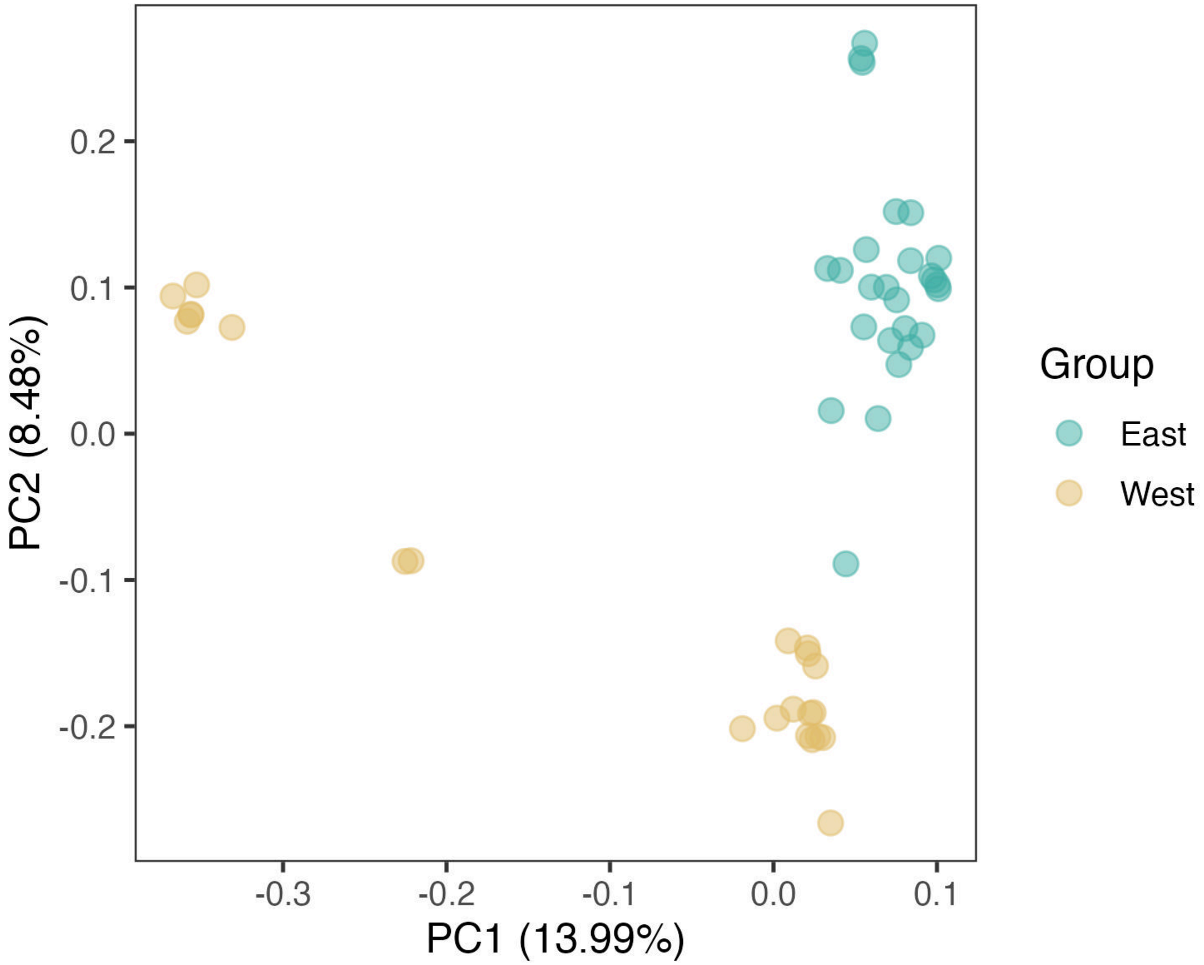


**Figure S5: PCA on genotype likelihoods for the high coverage WGS data.** The PCA was run on 30,000 sites. The high coverage WGS dataset contained only 6x . Colors and population ID follow Fig. 2.

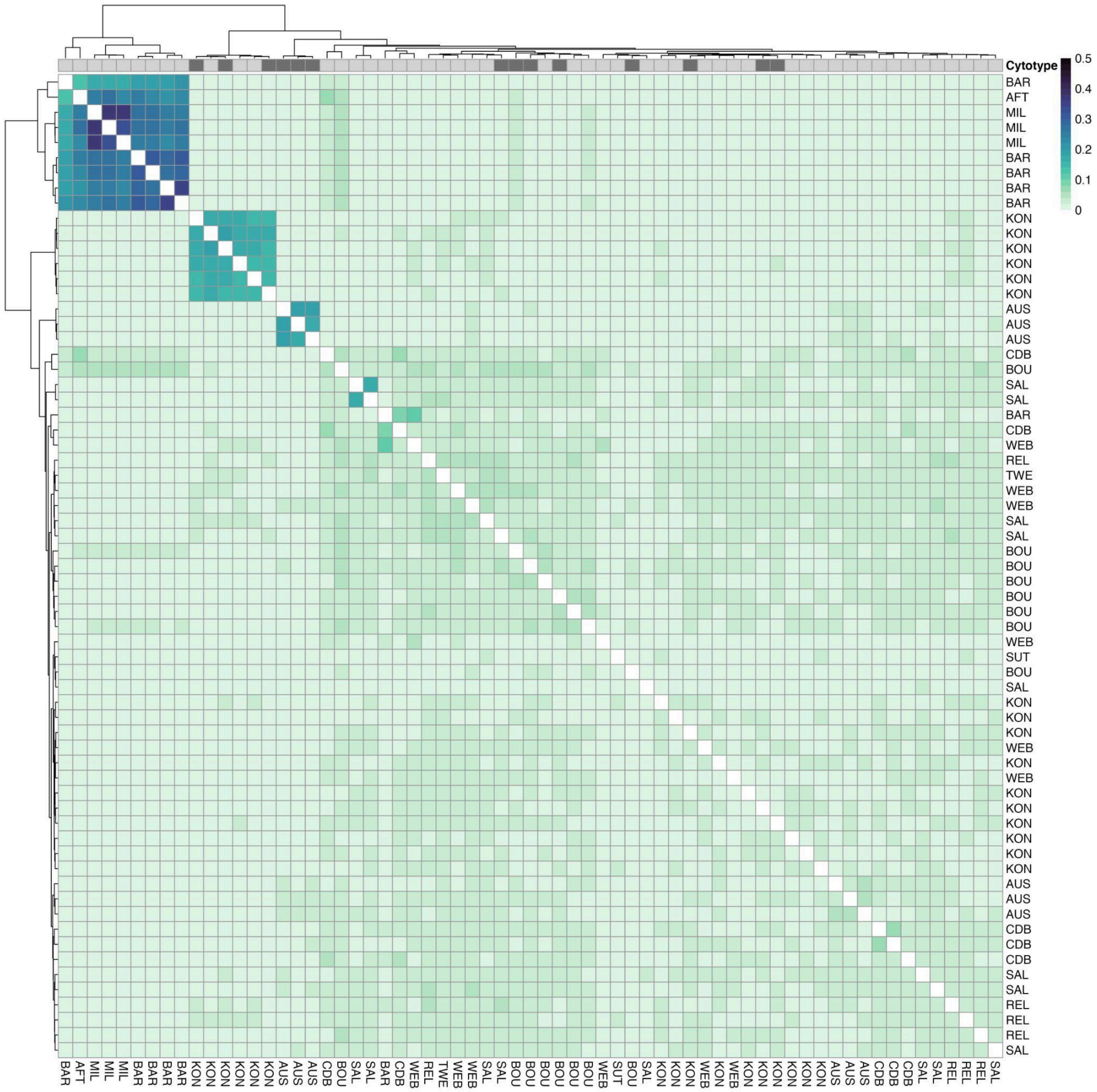


**Figure S6: Kinship of all sequenced genotypes in the West genetic group.** Columns are annotated with the cytotype of each genotype where 9*x* are dark gray and 6x are light gray. Genotypes are hierarchically clustered by Euclidean distance in kinship values and have the same order on both axes. Genotypes are labeled with their population code which follows Figure 2A. No data is plotted for the diagonal as estimates of self-relatedness are unreliable with low-coverage data.

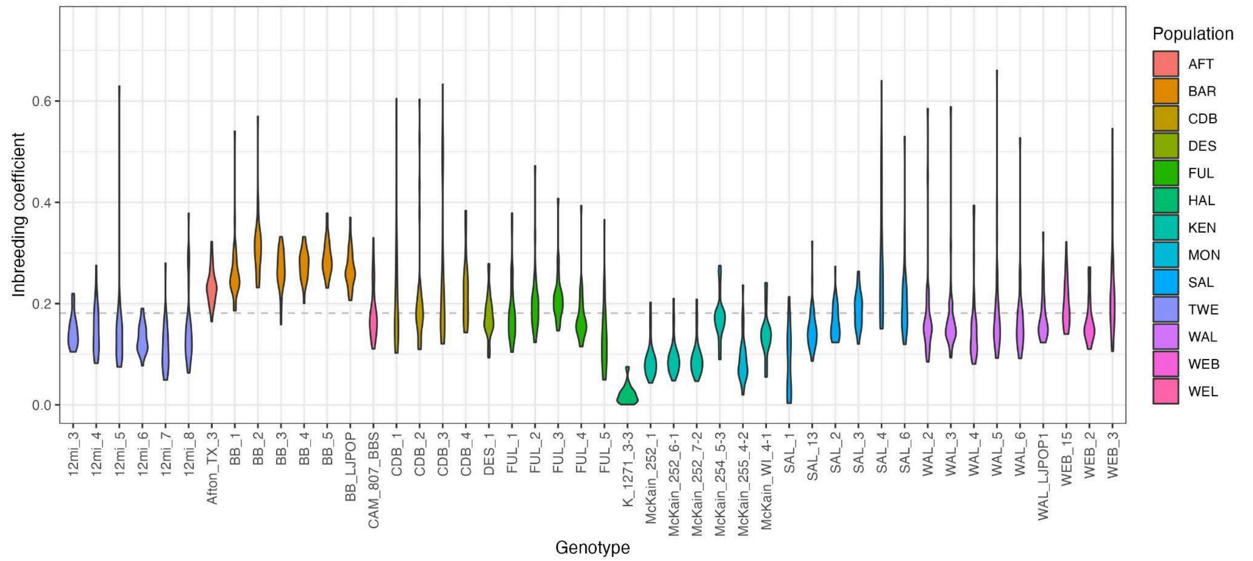


**Figure S7: Individual inbreeding coefficients estimated for hexaploid genotypes.** The violin plots show the distribution of the inbreeding coefficients estimated for each chromosome (n = 30) per genotype and are colored by population. The gray dashed line is the average inbreeding coefficient across these genotypes. BAR had a significantly higher average inbreeding coefficient than other populations ($F_{BAR}$ = 0.28±0.01, $p < 0.05$; $\overline{F} = 0.18$).

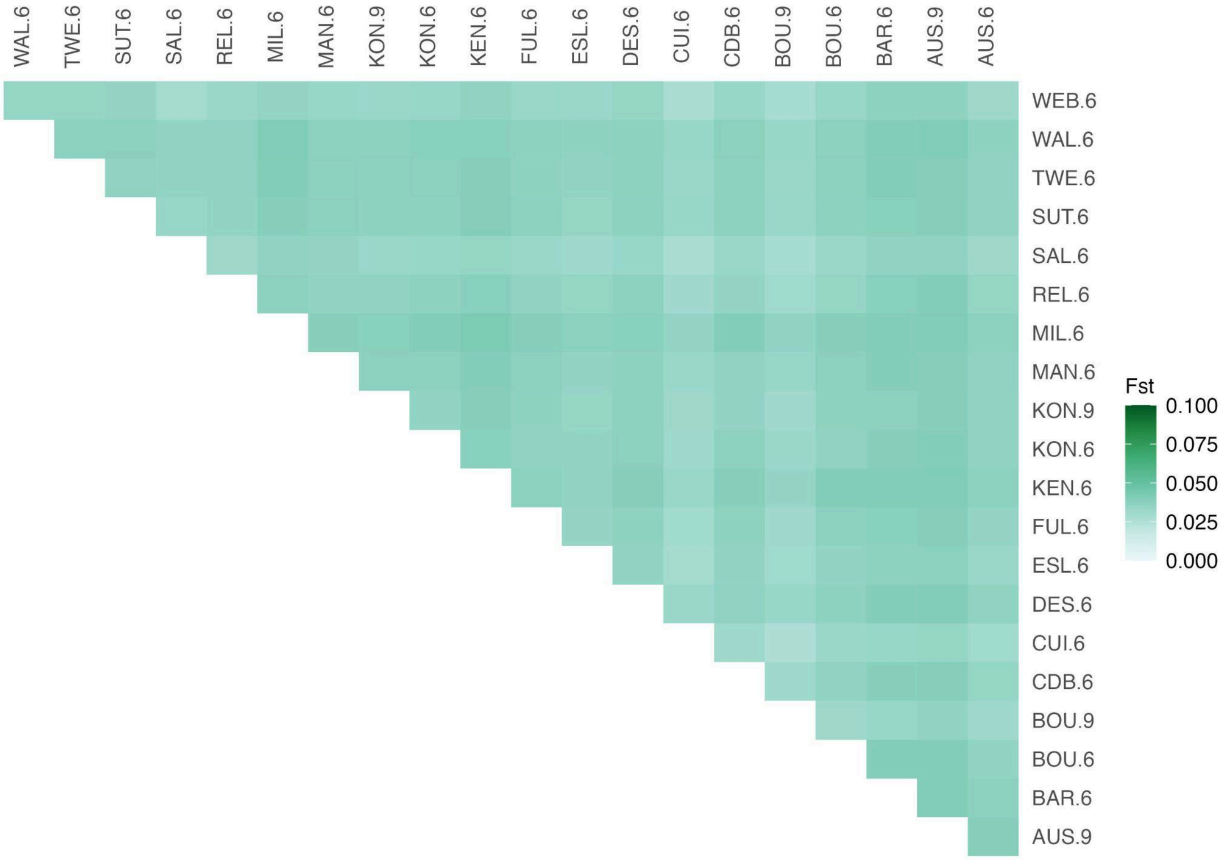


**Figure S8: Pairwise $F_{ST}$ estimated between 6*x* and 9*x* population pairs.** $F_{ST}$ is dependent on ploidy level and is expected to be elevated in comparisons among 9*x* compared to comparisons among 6*x* *(1)*. Only populations with greater than 3 genotypes were included.

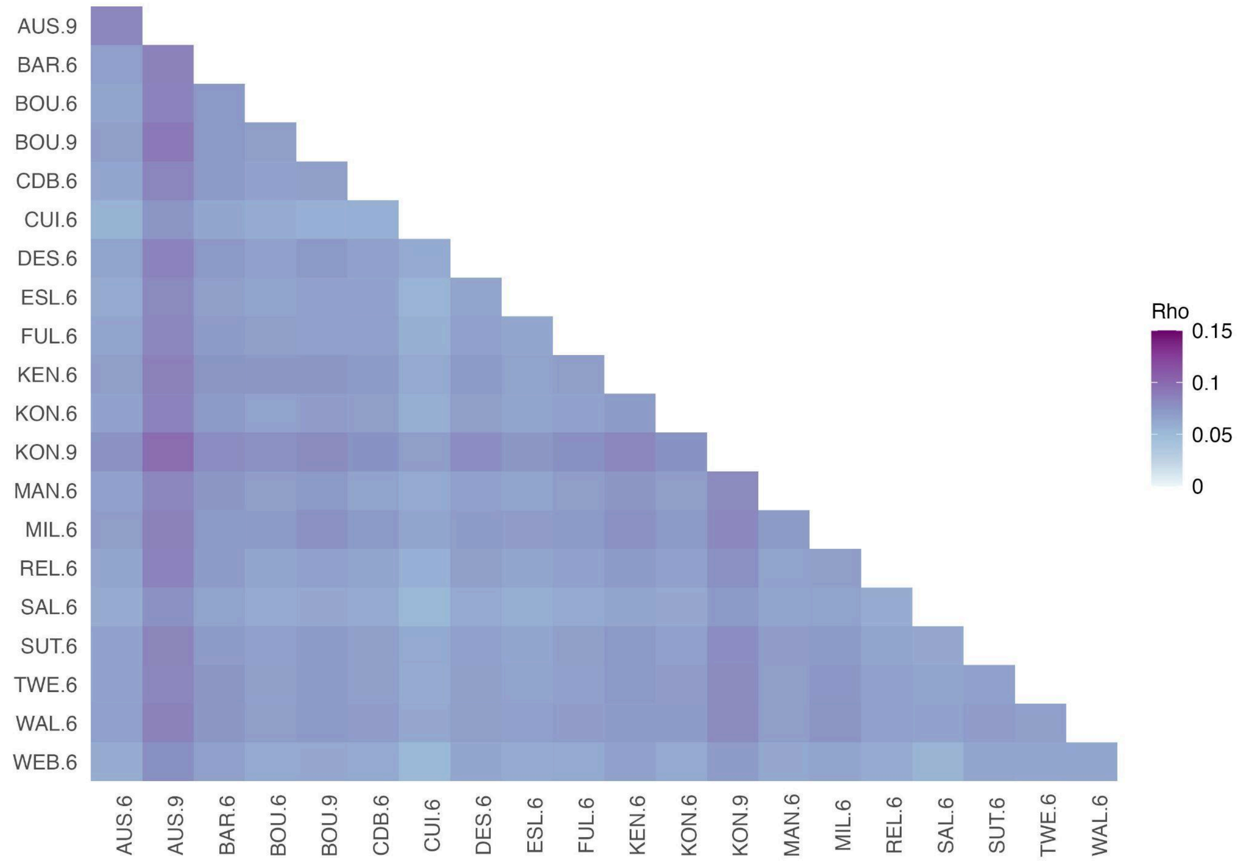


**Figure S9: Pairwise ρ estimated between 6*x* and 9*x* population pairs.** Only populations with greater than 3 genotypes were included.

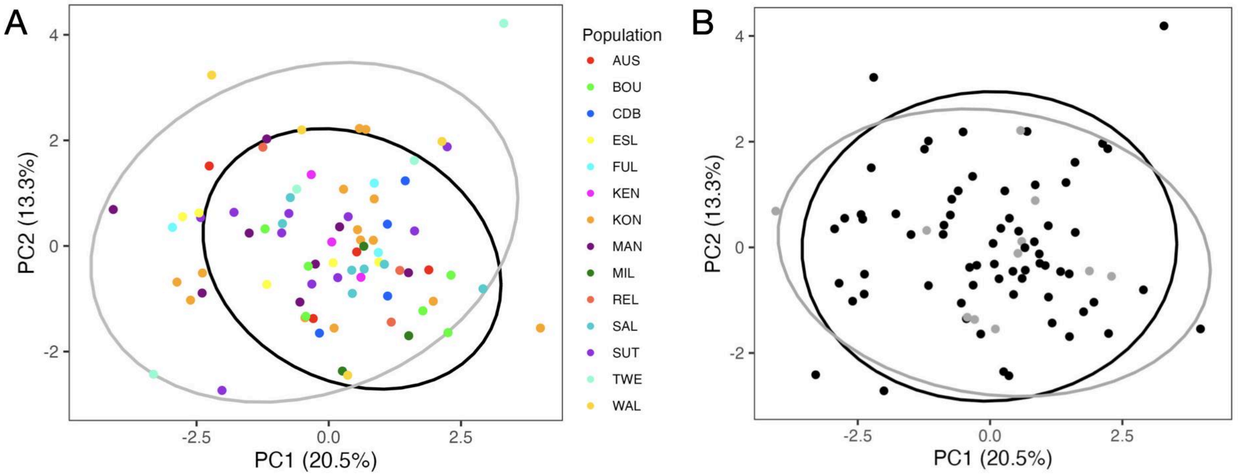


**Figure S11: PCA of phenotype BLUPs.** The first two principal components are plotted for each genotype overlain with the vectors for each standardized phenotype and 90% confidence ellipses. **(A)** Points are colored by population and the confidence ellipses enclose the East (black) andWest (gray) genetic groups. **(B)** Points and ellipses are colored by ploidy where the 6*x* cytotype is black and 9*x* cytotype is gray.

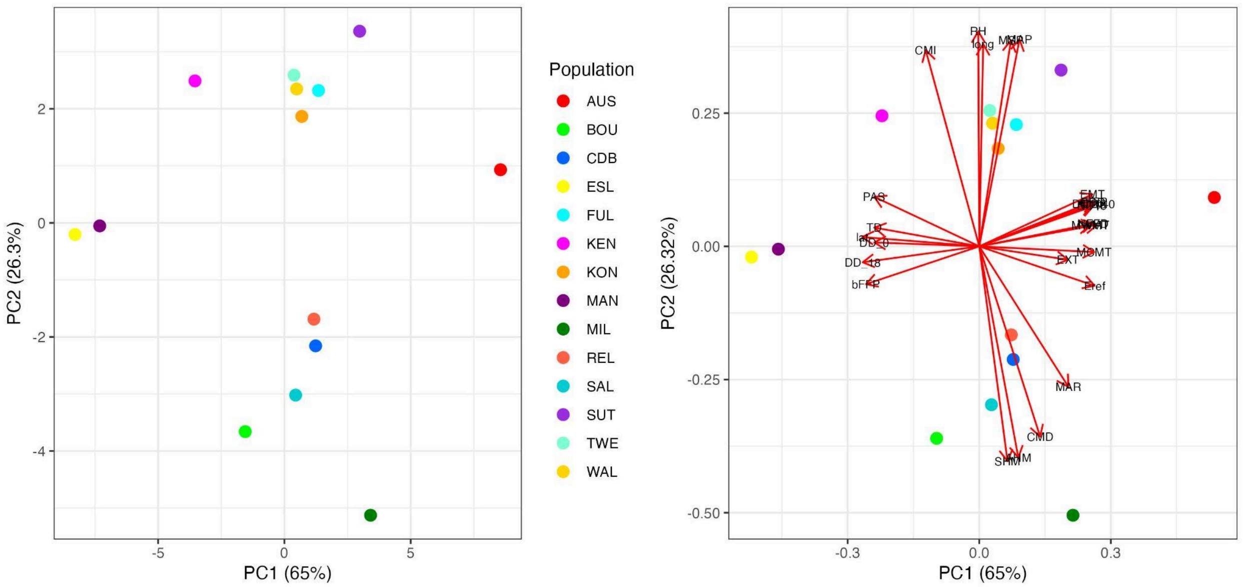


**Figure S12: Principal component loadings of the PCA on 28 climate variables. (Left)** PC1 and PC2 describe the majority of sampled environmental variation. **(Right)** The PC loadings overlaid as a biplot show high correlation among environmental variables. All climate variables are from ClimateNA except GDD, which is the mean yearly growing degree days at 10°C estimated with daymetr, latitude (lat), and longitude (lon). The climateNA variables are as follows: MAT = mean annual temperature (°C), MWMT = mean warmest month temperature (°C), MCMT = mean coldest month temperature (°C), TD = temperature difference between MWMT and MCMT, or continentality (°C), MAP = mean annual precipitation (mm), MSP = mean annual summer (May to Sept.) precipitation (mm), AHM = annual heat-moisture index (MAT+10)/(MAP1000)), SHM = summer heat-moisture index (MWMT)/(MSP1000), DD 0 = degree-days below 0◦C, DD5 = degree-days above 5°C, DD 18 = degree-days below 18°C, DD18 = degree-days above 18°C, NFFD = the number of frost-free days, FFP = frost-free period, bFFP = the day of the year on which FFP begins, eFFP = the day of the year on which FFP ends, PAS = precipitation as snow (mm), EMT = extreme minimum temperature over 30 years (°C), EXT = extreme maximum temperature over 30 years (°C), Eref = Hargreaves reference evaporation (mm), CMD = Hargreaves climatic moisture deficit (mm), MAR = mean annual solar radiation (MJ $m^{-2}$ $d^{-1}$), RH = mean annual relative humidity (%), CMI = Hogg's climate moisture index (mm), DD1040 = degree-days above 10°C and below 40°C.

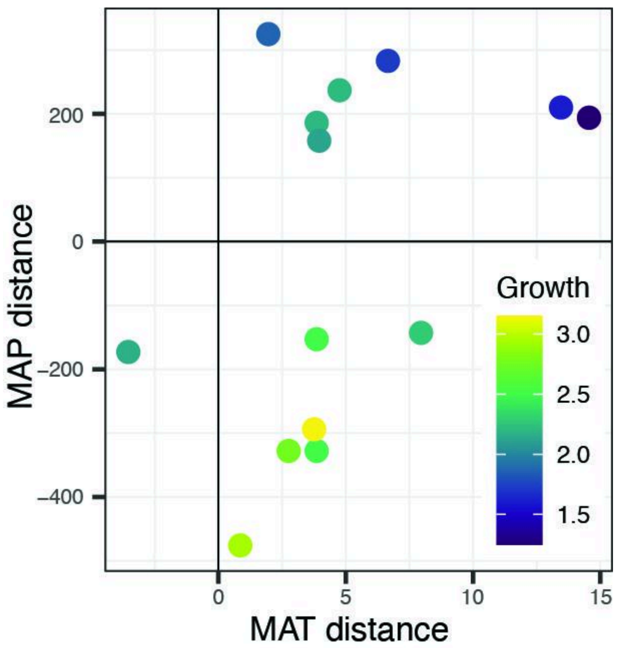


**Figure S13: Relative growth by MAP and and MAT distance.** Difference between the mean common garden environment and home environment of each population (garden - home) in mean annual precipitation (MAP) and mean annual temperature (MAT). The values for each population are plotted and colored by the population mean change in aboveground biomass. Populations in the lower right quadrant from colder and wetter environments performed the best while populations in the upper right quadrant, from colder and drier environments, performed the worst.

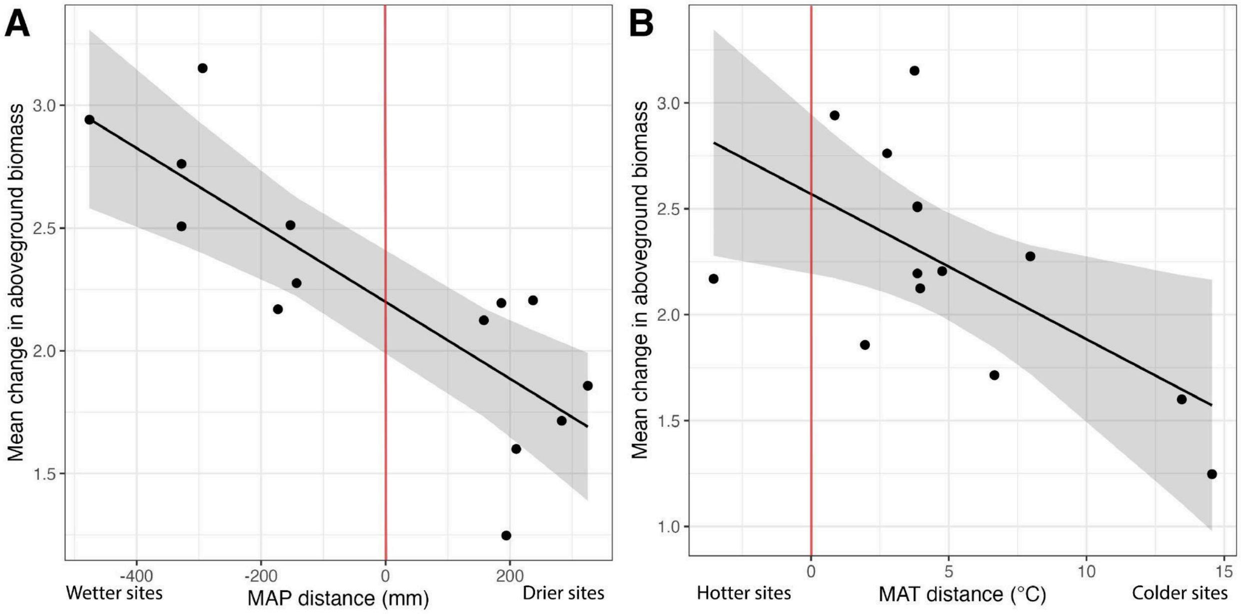


**Figure S14: Population mean change in aboveground biomass regressed against climate transfer**
**distance.** Predicted values (black line) are plotted with 95% confidence intervals in gray and population BLUPs overlaid as black dots. Climate transfer distance was estimated as (A) The difference in mean annual precipitation (MAP) and (B) mean annual temperature (MAT) between the common garden environment (averaged across 2021 and 2022) and the average home environment. The red line marks where the average common garden environment would be the same as a population's home environment.

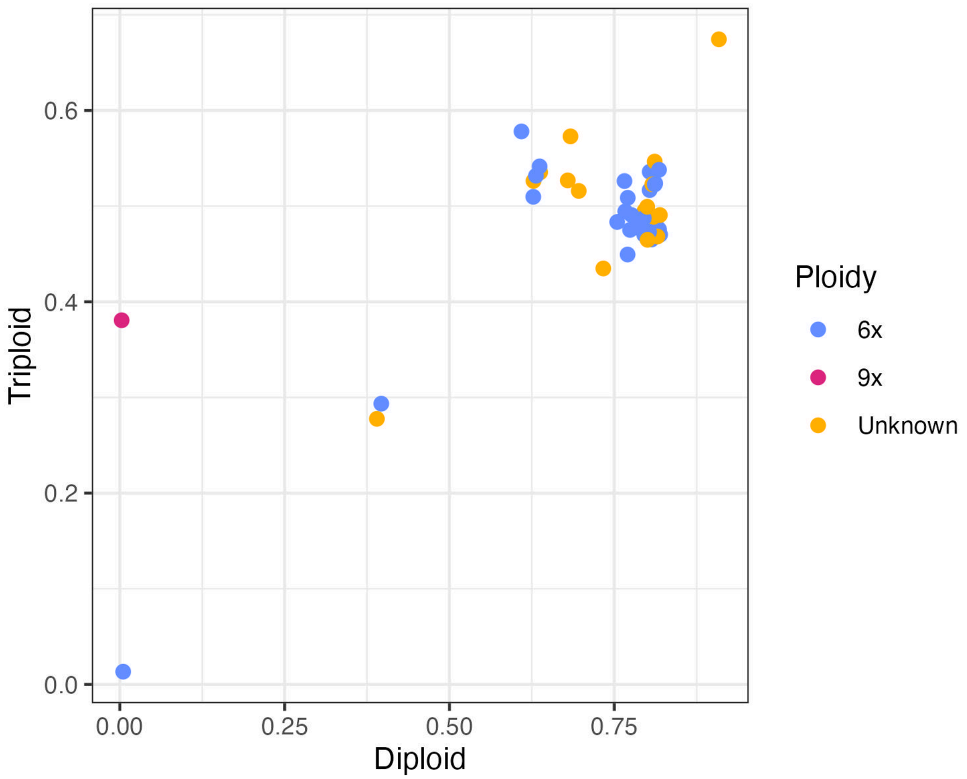


**Figure S15: The normalized maximized log-likelihood of the diploid and triploid nQuire models for genotypes sequenced to high coverage.** The color of the points represents whether the ploidy is unknown or known via flow cytometry. Of the genotypes for which ploidy was known, all were 6*x* except for one genotype. Tested unknown genotypes had similar values to known 6*x* genotypes except for two outliers. In the upper left and lower right corners are genotypes with relatively high (45x) and low (16x) coverage, respectively.

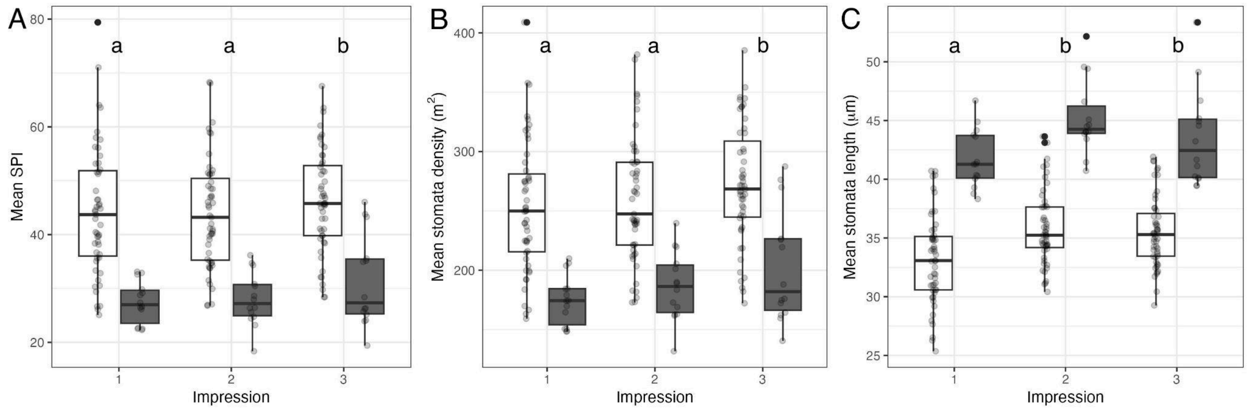


**Figure S16: The effect of cytotype on stomatal traits is consistent across the leaf.** Impression 1 was taken within the bottom 1 inch of the leaf, impression 2 was in the middle of the leaf, and impression 3 was within the top inch of the leaf. We assessed developmental variation in **(A)** SPI, **(B)** stomatal density, and **(C)** stomata length on a subset of genotypes (n = 23). The boxplot depict the distribution of genotype means for each ploidy where white is 6*x* and gray is 9*x* genotypes. The mean values for each genotype are overlaid as points. Lowercase letters above the boxplots indicate which impressions means are significantly different using a Tukey test and 95% confidence level.

### Supplemental Tables

**Table S1:** Genomic libraries included in the *Andropogon gerardi* (var. *Kellogg-1272*) genome assembly and their respective assembled sequence coverage levels in the final release. *Average read length of PacBio reads.

| **Library** | **Sequencing Platform** | **Average Read/Insert Size** | **Read Number** | **Assembled Sequence Coverage (x)** |
|---|---|---|---|---|
| JEXY | Illumina | 400 | 1,601,302,817 | 92.80 |
| GOXCG | Illumina-HiC | N/A | 938,370,083 | 54.38 |
| | PACBIO | 19,680* | 16,785,606 | 85.93 |
| **Total** | | N/A | 2,556,458,506 | 233.11 |

**Table S2:** PacBio library statistics for the libraries included in the *Andropogon gerardi* (var. *Kellogg-1272*) genome assembly and their respective assembled sequence coverage levels.

| Cutoff | Number of Reads | Basepairs | Average Read Length | Coverage |
|---|---|---|---|---|
| 0 | 16,785,606 | 343,726,563,435 | 19,680 | 85.93x |
| 1,000 | 16,784,941 | 343,726,421,728 | 19,680 | 85.93x |
| 2,000 | 16,782,985 | 343,723,282,795 | 19,681 | 85.93x |
| 3,000 | 16,781,398 | 343,719,397,953 | 19,681 | 85.93x |
| 4,000 | 16,780,091 | 343,714,820,784 | 19,682 | 85.93x |
| 5,000 | 16,777,997 | 343,705,307,473 | 19,682 | 85.93x |
| 6,000 | 16,774,286 | 343,684,717,768 | 19,683 | 85.92x |
| 7,000 | 16,768,962 | 343,650,000,372 | 19,684 | 85.91x |
| 8,000 | 16,763,136 | 343,606,292,932 | 19,686 | 85.90x |
| 9,000 | 16,757,799 | 343,560,982,110 | 19,687 | 85.89x |
| 10,000 | 16,753,026 | 343,515,663,833 | 19,688 | 85.88x |
| 11,000 | 16,748,251 | 343,465,530,496 | 19,689 | 85.87x |
| 12,000 | 16,743,169 | 343,406,976,519 | 19,690 | 85.85x |
| 13,000 | 16,734,980 | 343,303,965,973 | 19,692 | 85.83x |
| 14,000 | 16,704,343 | 342,886,573,490 | 19,699 | 85.72x |
| 15,000 | 16,566,666 | 340,874,186,260 | 19,732 | 85.22x |
| 16,000 | 16,066,795 | 333,079,495,791 | 19,853 | 83.27x |
| 17,000 | 14,753,317 | 311,321,403,055 | 20,189 | 77.83x |
| 18,000 | 12,463,595 | 271,193,752,445 | 20,850 | 67.80x |
| 19,000 | 9,938,051 | 224,485,620,806 | 21,750 | 56.12x |

**Table S3:** Summary statistics of the initial output of the HAP1 RACON polished HiFiAsm+HIC assembly. The table shows total contigs and total assembled basepairs for each set of scaffolds greater than the size listed in the left hand column.

| Minimum Scaffold Length | Number of Scaffolds | Number of Contigs | Scaffold Size | Basepairs | % Non-gap Basepairs |
|---|---|---|---|---|---|
| 5 Mb | 61 | 61 | 2,664,415,677 | 2,664,415,677 | 100.00% |
| 2.5 Mb | 65 | 65 | 2,677,111,928 | 2,677,111,928 | 100.00% |
| 1 Mb | 68 | 68 | 2,681,267,982 | 2,681,267,982 | 100.00% |
| 500 Kb | 77 | 77 | 2,687,222,053 | 2,687,222,053 | 100.00% |
| 250 Kb | 117 | 117 | 2,700,371,179 | 2,700,371,179 | 100.00% |
| 100 Kb | 344 | 344 | 2,732,853,625 | 2,732,853,625 | 100.00% |
| 50 Kb | 1,064 | 1,064 | 2,780,484,710 | 2,780,484,710 | 100.00% |
| 25 Kb | 1,064 | 1,064 | 2,780,484,710 | 2,780,484,710 | 100.00% |
| 10 Kb | 1,064 | 1,064 | 2,780,484,710 | 2,780,484,710 | 100.00% |
| 5 Kb | 1,064 | 1,064 | 2,780,484,710 | 2,780,484,710 | 100.00% |
| 2.5 Kb | 1,064 | 1,064 | 2,780,484,710 | 2,780,484,710 | 100.00% |
| 1 Kb | 1,064 | 1,064 | 2,780,484,710 | 2,780,484,710 | 100.00% |
| 0 bp | 1,064 | 1,064 | 2,780,484,710 | 2,780,484,710 | 100.00% |

**Table S4:** Summary statistics of the initial output of the HAP2 RACON polished HiFiAsm+HIC assembly. The table shows the total contigs and total assembled basepairs for each set of scaffolds greater than the size listed in the left-hand column.

| Minimum Scaffold Length | Number of Scaffolds | Number of Contigs | Scaffold Size | Basepairs | % Non-gap Basepairs |
|---|---|---|---|---|---|
| 5 Mb | 61 | 61 | 2,608,260,179 | 2,608,260,179 | 100.00% |
| 2.5 Mb | 65 | 65 | 2,622,080,605 | 2,622,080,605 | 100.00% |
| 1 Mb | 67 | 67 | 2,625,544,079 | 2,625,544,079 | 100.00% |
| 500 Kb | 78 | 78 | 2,633,098,987 | 2,633,098,987 | 100.00% |
| 250 Kb | 115 | 115 | 2,645,787,790 | 2,645,787,790 | 100.00% |
| 100 Kb | 290 | 290 | 2,671,355,876 | 2,671,355,876 | 100.00% |
| 50 Kb | 731 | 731 | 2,701,527,046 | 2,701,527,046 | 100.00% |
| 25 Kb | 731 | 731 | 2,701,527,046 | 2,701,527,046 | 100.00% |
| 10 Kb | 731 | 731 | 2,701,527,046 | 2,701,527,046 | 100.00% |
| 5 Kb | 731 | 731 | 2,701,527,046 | 2,701,527,046 | 100.00% |
| 2.5 Kb | 731 | 731 | 2,701,527,046 | 2,701,527,046 | 100.00% |
| 1 Kb | 731 | 731 | 2,701,527,046 | 2,701,527,046 | 100.00% |
| 0 bp | 731 | 731 | 2,701,527,046 | 2,701,527,046 | 100.00% |

**Table S5:** Final summary assembly statistics for the HAP1 chromosome scale assembly.

| | |
|---|---|
| **Scaffold total** | 48 |
| **Contig total** | 88 |
| **Scaffold sequence total** | 2,669.7 Mb |
| **Chromosome Sequence** | 2,666.8 Mb |
| **Contig sequence total** | 2,669.3 Mb (0.001% gap) |
| **Scaffold N/L50** | 13 / 86.8 Mb |
| **Contig N/L50** | 16 / 63.1 Mb |

**Table S6:** Final summary assembly statistics for the HAP2 chromosome scale assembly.

| | |
|---|---|
| **Scaffold total** | 39 |
| **Contig total** | 68 |
| **Scaffold sequence total** | 2,588.6 Mb |
| **Chromosome Sequence** | 2,586.5 Mb |
| **Contig sequence total** | 2,588.3 Mb (0.05% gap) |
| **Scaffold N/L50** | 13 / 86.4 Mb |
| **Contig N/L50** | 17 / 59.2 Mb |

### Supporting Text

### 1. Supplemental Methods

#### 1.1 Identifying sequencing contamination

To assess the quality of the short-read sequences from Cornell and detect possible contamination, reads from each sample were aligned to the sorghum reference genome (NCBI GenBank ID GCF_000003195.3) using bwa mem (2), as the A. gerardi reference genome was still in assembly. Alignment statistics included the fraction of mapped reads, duplication rate, the fraction of bases of the whole genome and of the coding sequence portion covered at multiple depths (>0x, >1x, >5x), and the fraction of reads mapping with various numbers of mismatches (0, ≤5, ≤10, ≤15). The Kraken pipeline (3) was used to quantify contamination for each sample with sequences originating from bacteria, the human genome, and plants outside of Poaceae. To further confirm the taxonomy of the analyzed sequences, a custom database of five plastid genes (*matK, ndhF, rbcL, rpoB* and *rpoC1*) was constructed from 4,755 plant plastid genomes downloaded from NCBI RefSeq (Dataset 01), all of which  have species-level taxonomic information. Nucleotide sequences of the five genes were extracted from the plastid genomes of each species based on NCBI Refseq gene annotation. For genomes without gene annotations, TBLASTN was used to identify the coordinates of these genes within the respective genomes. For each sample, bwa aln (4) was then used to map 1,000 randomly selected reads to the plastid genes database. Up to five such genes with the most reported hits were selected, and the Phylum, Class, Order, Family, Subfamily, Tribe, Genus, and Species of these genes were reported. While the categories Phylum through Tribe were typically as expected (Streptophyta, Magnoliopsida, Poales, Poaceae, Panicoideae, Andropogoneae, respectively), Genus was often ambiguous between *Schizachyrium* and *Andropogon* due to the species' allopolyploidy. 9 samples (6%) were identified as anything other than *Schizachyrium* or *Andropogon* and were discarded.

#### 1.2 Resolving genome contamination and error correction

Additional scaffolds were classified in HAP1 as repetitive (>95% masked with 24-mers that occur more than 4 times in the chromosomes; 786 scaffolds, 87.8 Mb), redundant (unanchored scaffolds composed of ≥95% 24-mers >2x in all scaffolds; 2 scaffolds, 31.1 kb), mitochondria (177 scaffolds, 10.5 Mb), and prokaryote (13 scaffolds, 727.0 kb). Scaffolds were also classified in HAP2 as repetitive (>95% masked with 24-mers that occur more than 4 times in the chromosomes; 554 scaffolds, 66.8 Mb), redundant (unanchored scaffolds composed of ≥95% 24-mers >2x in all scaffolds; 5 scaffolds, 115.9 kb), and mitochondria (83 scaffolds, 5.1 Mb).

After forming the chromosomes, it was observed that some small (<20 kb) redundant sequences were present on adjacent contig ends within chromosomes. To resolve this issue, adjacent contig ends were aligned to one another using BLAT (v35, 5), and duplicate sequences were collapsed to close the gap between them. A total of 2 adjacent contig pairs were collapsed in the HAP1 assembly and 6 in the HAP2 assembly.

Heterozygous SNP/indel phasing errors were corrected using the 85.93x CCS data. A total of 1,903 heterozygous SNPs/indels were corrected in both haplotypes. Homozygous SNPs and indels were corrected in both haplotypes using ~62x of Illumina reads (2x150, 400 bp insert) by aligning the reads using bwa mem (v0.7.17-r1188, 2) and identifying homozygous SNPs and indels with the GATK's UnifiedGenotyper tool (v3.6-0-g89b7209, 6). A total of 987 homozygous SNPs and 10,691 homozygous indels were corrected in HAP1 and 882 homozygous SNPs and 9,487 homozygous indels were corrected in HAP2.

#### 1.3 PASA gene models

Loci were determined by transcript assembly alignments, EXONERATE alignments, and Swiss-Prot proteomes to identify repeats and soft-masked *A. gerardi* respective genomes using RepeatMasker (7) with up to 2,000 bp extension on both ends unless extending into another locus on the same strand. EXONERATE alignments used protein sequences from *Arabidopsis thaliana*, *Glycine max*, *Oryza sativa*, *Sorghum bicolor*, *Brachypodium distachyon*, *Aquilegia coerulea*, *Solanum lycopersicum*, *Vitis vinifera*, *Panicum hallii*, *Joinvillea ascendens*, *Acorus americanus*, *Paspalum vaginatum*, *Phoenix dactylifera*, *Musa acuminata*, *Ananas comosus*, *Asparagus officinalis*, and *Phalaenopsis equestris*. The repeat library consists of *de novo* repeats by RepeatModeler on *A. gerardi* HAP1 and repeats in RepBase (8). Gene models were predicted by homology-based predictors, FGENESH+ (9), FGENESH EST (similar to FGENESH+, but using EST to compute splice site and intron input instead of protein/translated open reading frame, ORFs), EXONERATE (10), PASA assembly ORFs (a JGI homology constrained ORF finder), and AUGUSTUS (11) trained by the high confidence PASA assembly ORFs and with intron hints from short read alignments. The best-scored predictions for each locus were selected using multiple positive factors, including EST and protein support, and one negative factor of overlap with repeats. PASA improved the selected gene predictions by adding untranslated regions, splicing correction, and alternative transcripts.

### 2. Extended analysis of phenotype data

Here, we provide a description of the measured phenotypes we found to be unaffected by ploidy. We report and discuss the direct effects of environmental PC1 and PC2 on trait variation while controlling for ploidy, and how this informs our understanding of the leaf economic spectra in *A. gerardi*. The discussion is structured by grouping leaf morphology traits and performance traits. Stomatal traits and relative growth are not discussed as they are comprehensively addressed in the main text.

#### 2.1. Leaf morphology traits

Leaf morphology traits (width, length, thickness) were significantly genetically correlated in year 1; the direction of the relationships was consistent in year 2, although not significantly different from zero (Fig. 3C). Only leaf thickness was significantly genetically correlated with specific leaf area (SLA) and no leaf morphology traits were significantly correlated with leaf dry matter content (LDMC). These results suggest there is a constraint on overall leaf morphology where a limited dimension of leaf shape is available. For example, no *A. gerardi* individuals have leaves that are 10 cm wide and 4 cm long (broad and short). As precipitation (environmental PC2) increases, genotypes have wider and thinner leaves (Fig. S1, S4), consistent with a fast-slow growth strategy trade-off where genotypes in drier climates have a more conservative strategy (12). Interestingly, leaf size and leaf thickness increases with temperature and growing season length (environmental PC1; Fig. S1-4). This is consistent with selection for more robust leaves under drier climates.

Although there is constraint on overall leaf morphology, we found these traits have the lowest heritability of the measured traits in our experiment suggesting high phenotypic plasticity within the trait space (Fig. 3C). Phenotypic plasticity has been previously described in *A. gerardi* physiology, root architecture, and performance traits (13–17) and is a known drought tolerance mechanism (18). Within a single season, it has been demonstrated that an *A. gerardi* plant will utilize a drought response strategy of rapid leaf senescence and high leaf turnover, creating a canopy of younger leaves with lower SLA (lower mass per leaf area), increase resource allocation root biomass, and decrease allocation to flowering (19). Together, this suggests that although there are constraints on individual leaf traits, the overall leaf shape is quite plastic.

#### 2.1.1 Leaf width

We found leaf width was not significantly different between cytotypes (Model 1, Figure S1C). We did find a significant effect of environmental PC1 and PC2 (Model 2, Figure S1D-E); leaf width increases with precipitation and length of the growing season.

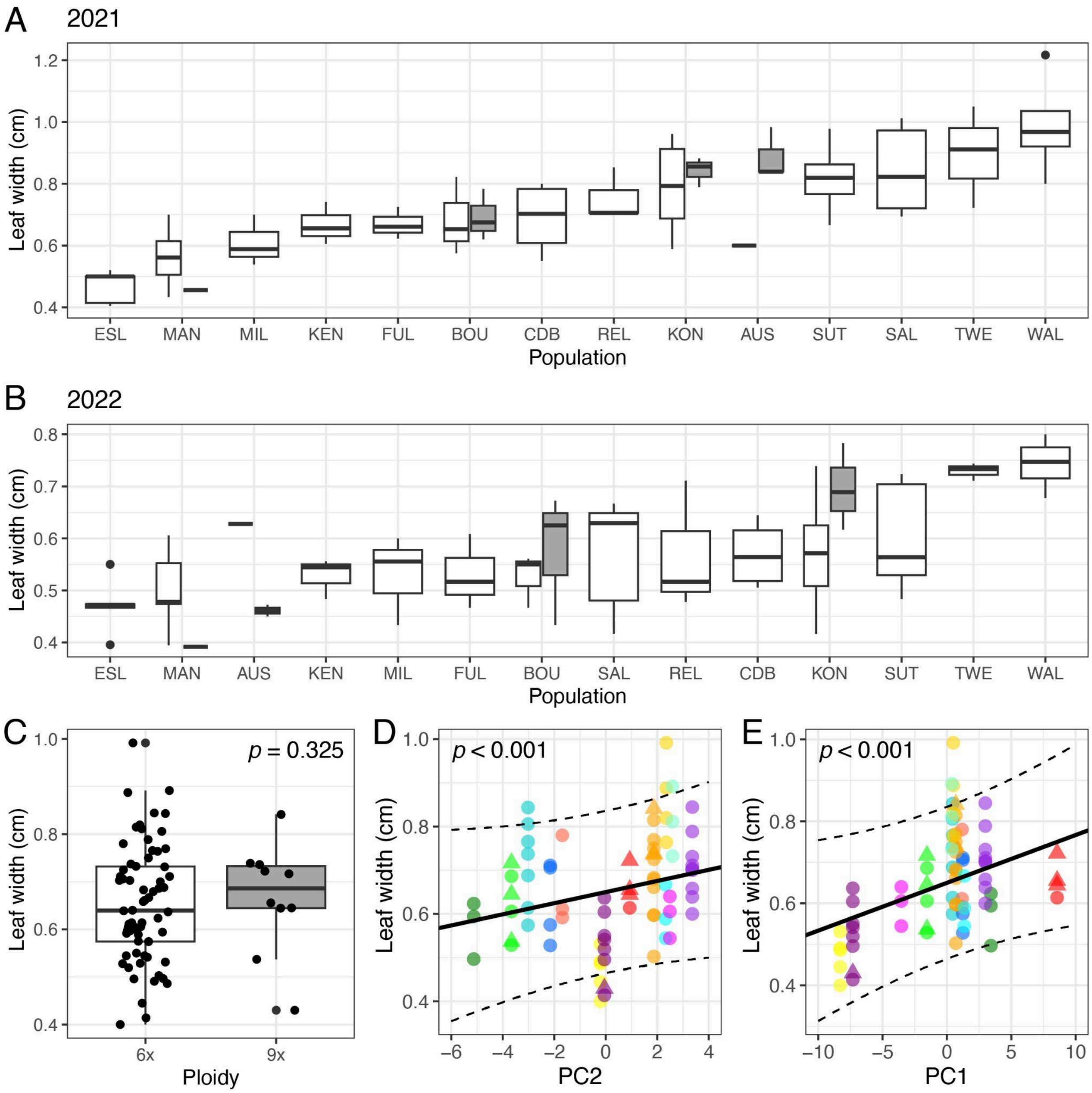


**Figure S17. Variation in leaf width. (A-B)** Population variation in leaf length ordered by population means. Data is plotted separately for each ploidy in mixed-ploidy populations (white = 6*x*, grey = 9*x*) and separately for the data collections years of 2021 and 2022. **(C)** Leaf width by ploidy, where black dots are the mean phenotype for each genotype. **(D-E)** Leaf width over environmental PC2 and PC1, respectively. Points are the mean phenotype for each genotype with color specifying population and shape specifying ploidy (following color and shape codes in Fig. 3). The black line is the predicted effect of the PC2 or PC1 on leaf width and the dashed lines are two standard errors from the mean. In all panels, a p-value less than 0.0033 is considered significant at a confidence level of 95%, after a Bonferroni correction. The p-value in C-E refers to the significance of the variable on the x-axis.

#### 2.1.2 Leaf length

We found the leaf length was not significantly affected by cytotype using Model 1 (Figure S2C). PC1 had a significant effect but PC2 did not (Model 2, Figure S2D-E). Leaf length increased with length of the growing season, similarly to leaf width.

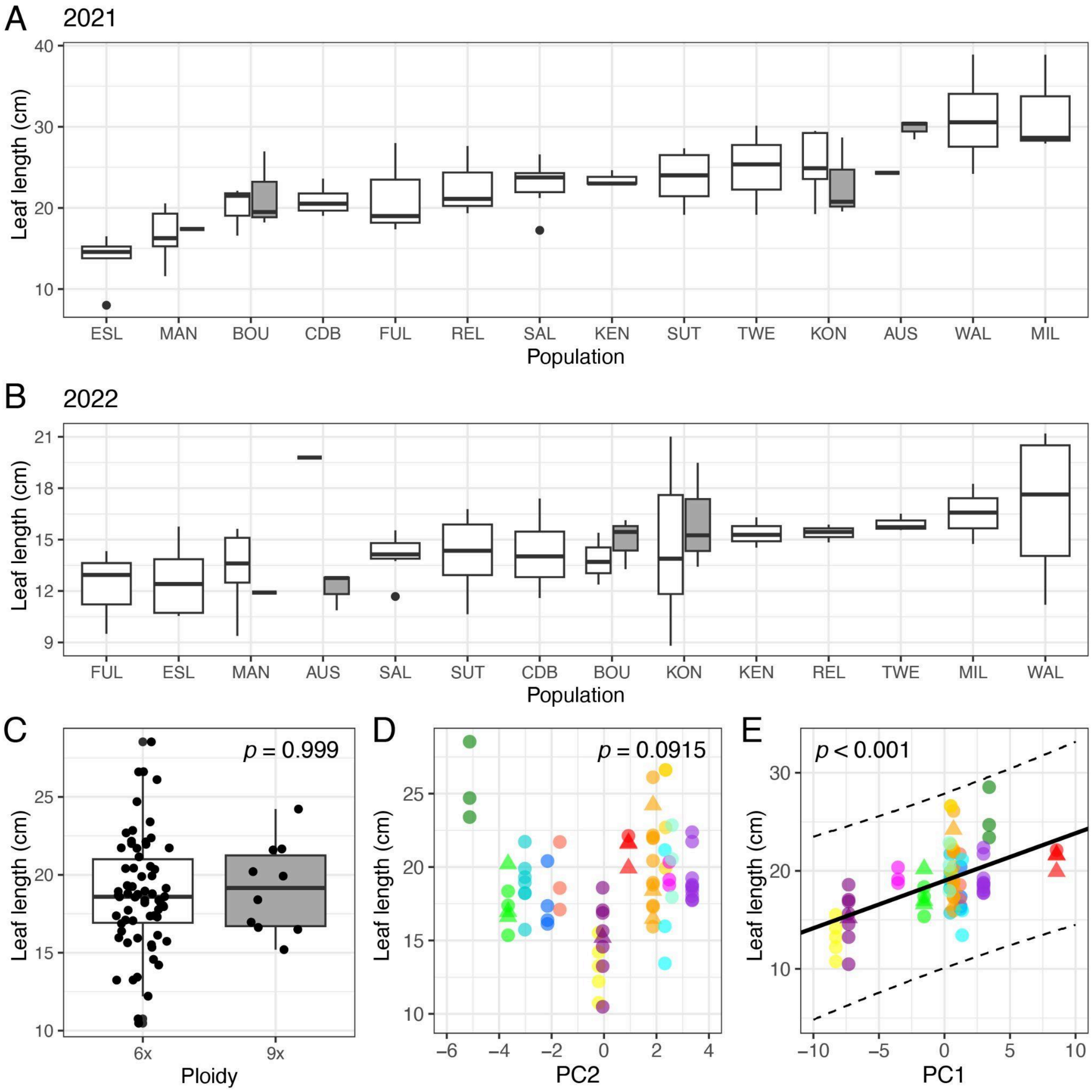


**Figure S18. Variation in leaf length.** See Fig. S1 for figure descriptions.

#### 2.1.3 Leaf thickness

As leaf thickness is strongly correlated with leaf length and width, we found cytotype did not have a significant effect (Model 1, Figure S3C). We only found a significant effect of PC1, and not PC2, on leaf thickness, where leaf thickness increases with length of the growing season (Model 2, Figure 3D-E).

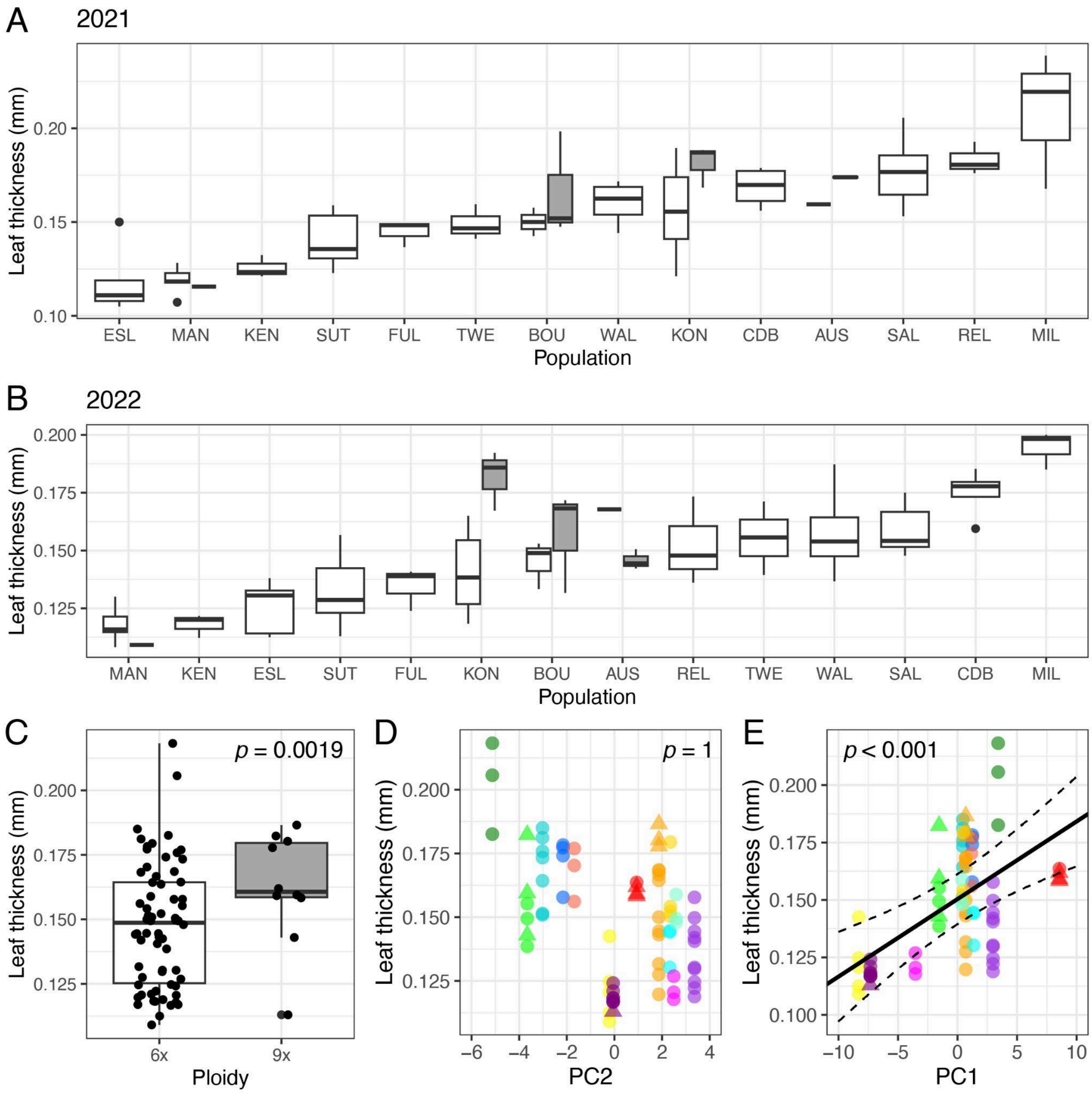


**Figure S19. Variation in leaf thickness.** See Fig. S1 for figure descriptions. No prediction line is plotted in D as the relationship is not significant.

#### 2.1.4 Specific leaf area

We found that specific leaf area (SLA) does not vary significantly between cytotypes (Model 1, Figure S4C). We saw a significant effect of both PC1 and PC2 on SLA, where SLA decreases with length of the growing season and increases with precipitation (Model 2, Figure 4D-E). This is opposite of the effect of PC1 on leaf thickness, which is consistent with SLA and leaf thickness being negatively genetically correlated (Fig. 2C). SLA is often regulated by water and carbon economics (20). A lower SLA indicates greater investment of resources in individual leaves (dry mass) and longer leaf longevity whereas a higher SLA can indicate a greater photosynthetic capacity and greater competitive ability in resource-rich environments (21). The positive relationship of SLA with PC2 may represent variation along the leaf economic spectrum; where *A. gerardi* genotypes have limited water and a low SLA, a more conservative growth strategy may be used (Fig. S4D), as opposed to a fast-growth strategy in an environment with ample water.

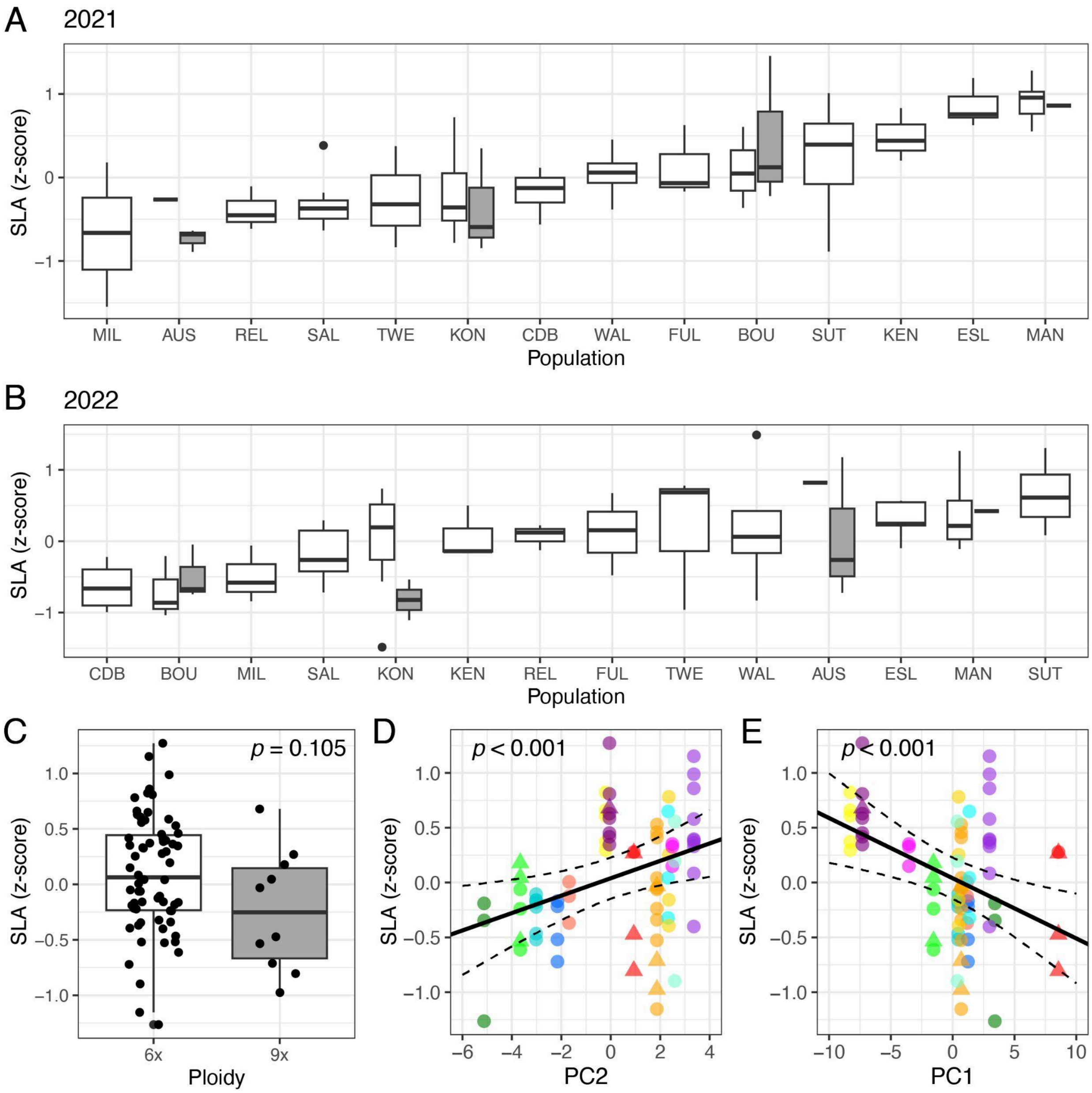


**Figure S20. Variation in SLA.** See Fig. S1 for figure descriptions. SLA was transformed using an inverse-normal transform. As such, the transformed values (Z-scores) are unitless.

#### 2.1.5 Leaf dry matter content

We found leaf dry matter content (LDMC) did not vary significantly between cytotypes (Model 1, Figure S5C), nor did either environmental PC (Model 2, Figure 5C-D). As SLA is inversely related to LDMC (20), the limited variation in LDMC variation suggests that the variation in SLA is largely a result of changes in leaf area, rather than leaf structure.

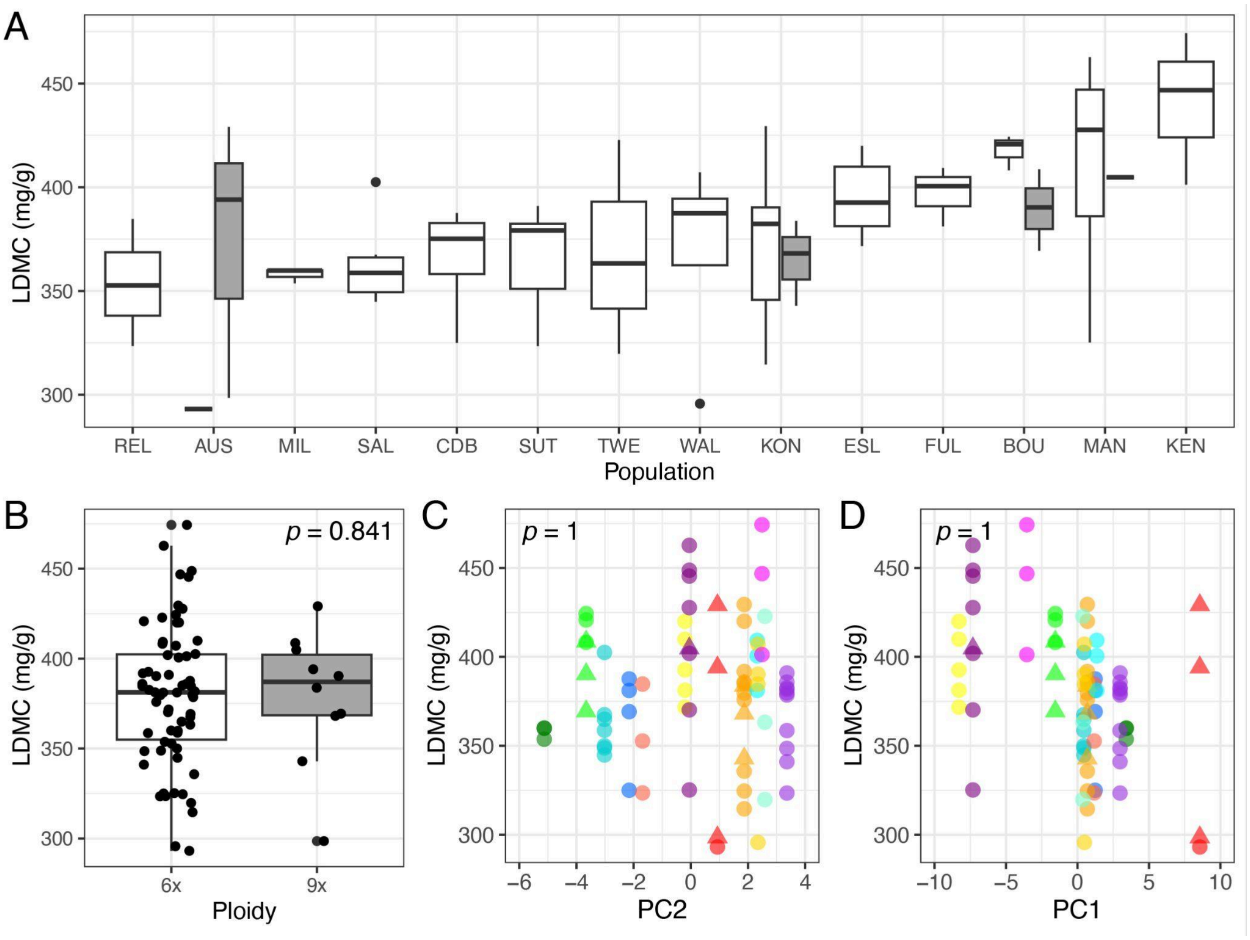


**Figure S21. Variation in LDMC. (A)** Population variation in LDMC ordered by population means. Data is plotted separately for each ploidy in mixed-ploidy populations (white = 6x, grey = 9x). **(B)** LDMC by ploidy, where black dots are the mean phenotype for each genotype. **(C-D)** LDMC over environmental PC2 and PC1, respectively. Points are the mean phenotype for each genotype with color specifying population and shape specifying ploidy (following color and shape codes in Fig. 3). In all panels, a p-value less than 0.0033 is considered significant at a confidence level of 95%, after a Bonferroni correction. The p-value in B-D refers to the significance of the variable on the x-axis. No predictive lines are plotted in C-D as the relationship is insignificant.

#### 2.2. Performance traits

The performance traits measured can provide different information through examining the absolute trait value versus the change between years. For example, plant height is expected to vary between populations due to pressure from historical grazing and climate regimes but a change in plant height between years can reflect growth rate or plant age class (16, 22). As such, we examined growth variables as well as the absolute trait values.

#### 2.2.1 Height

We found plant height was not affected by ploidy (Model 1, FIg. S6C) but was significantly associated with environmental PC1 and PC2 (Model 2, Fig. S6D-E). The shortest plants were those from the most Northern population (Fig. S6A-B) and the tallest were from mid-latitude populations with high precipitation and warm temperatures. Plant height had the highest heritability of the measured traits in this experiment (Fig. 3C). These results are consistent with previous studies demonstrating genetic control of plant height, with a strong effect of growing season length and precipitation (14, 23, 24).

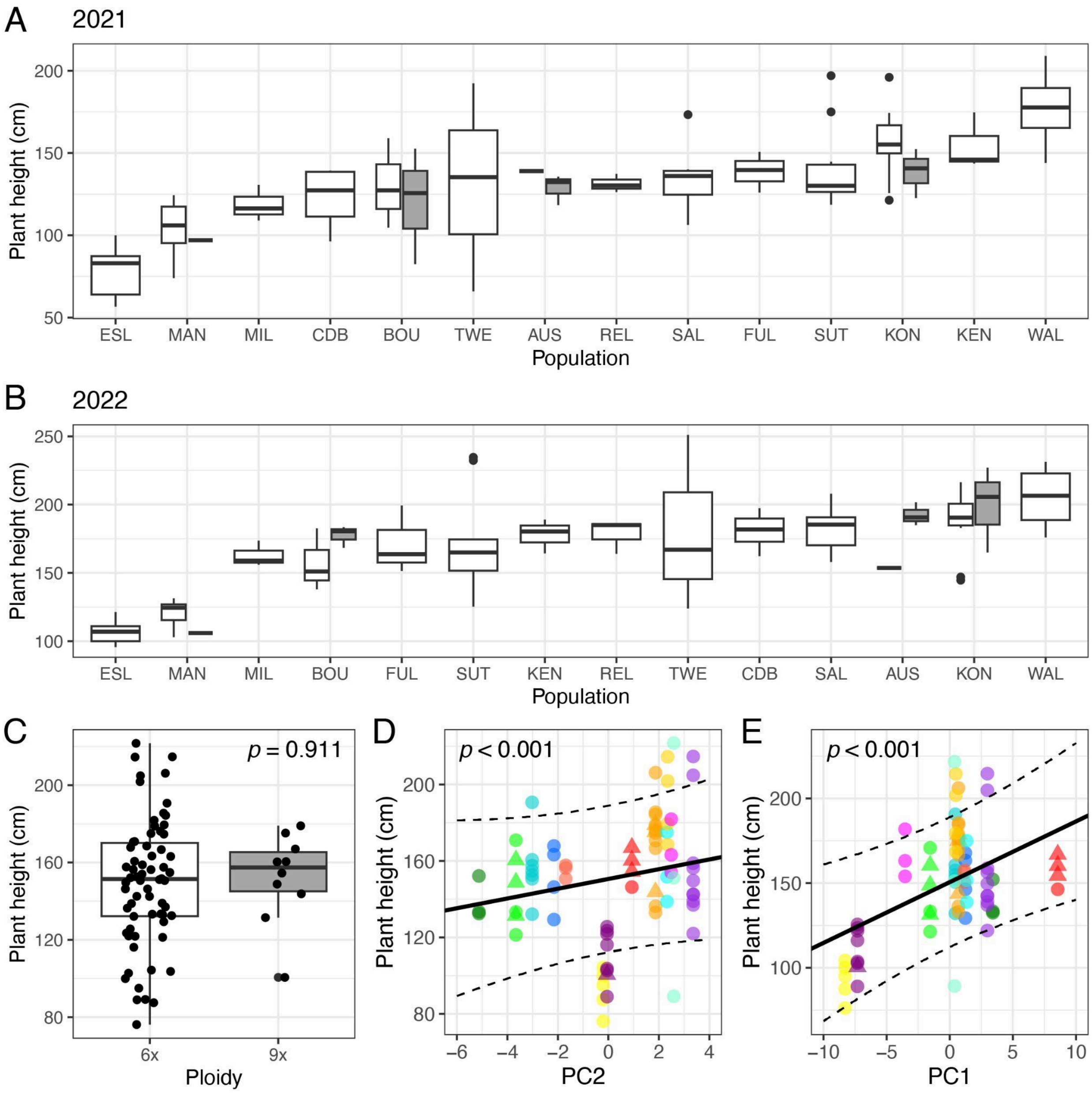


**Figure S22. Variation in plant height.** See Fig. S1 for figure descriptions.

#### 2.2.2 Change in plant height (height growth)

Change in plant height between years 1 and 2 (height growth) was not significantly affected by ploidy (Model 1, Fig. S7B) nor environmental PC2 (Model 2, Fig. S7C). Rather, variation in height growth was significantly affected by environmental PC1 (Model 2, Fig. S7D), where change in plant height increased with temperature and length of the growing season. If short plant height is selected for in climates with shorter growing seasons and colder temperatures (Fig. 6D), the degree of change in plant height is expected to be higher for those in longer growing seasons and warmer temperatures, as is seen here.

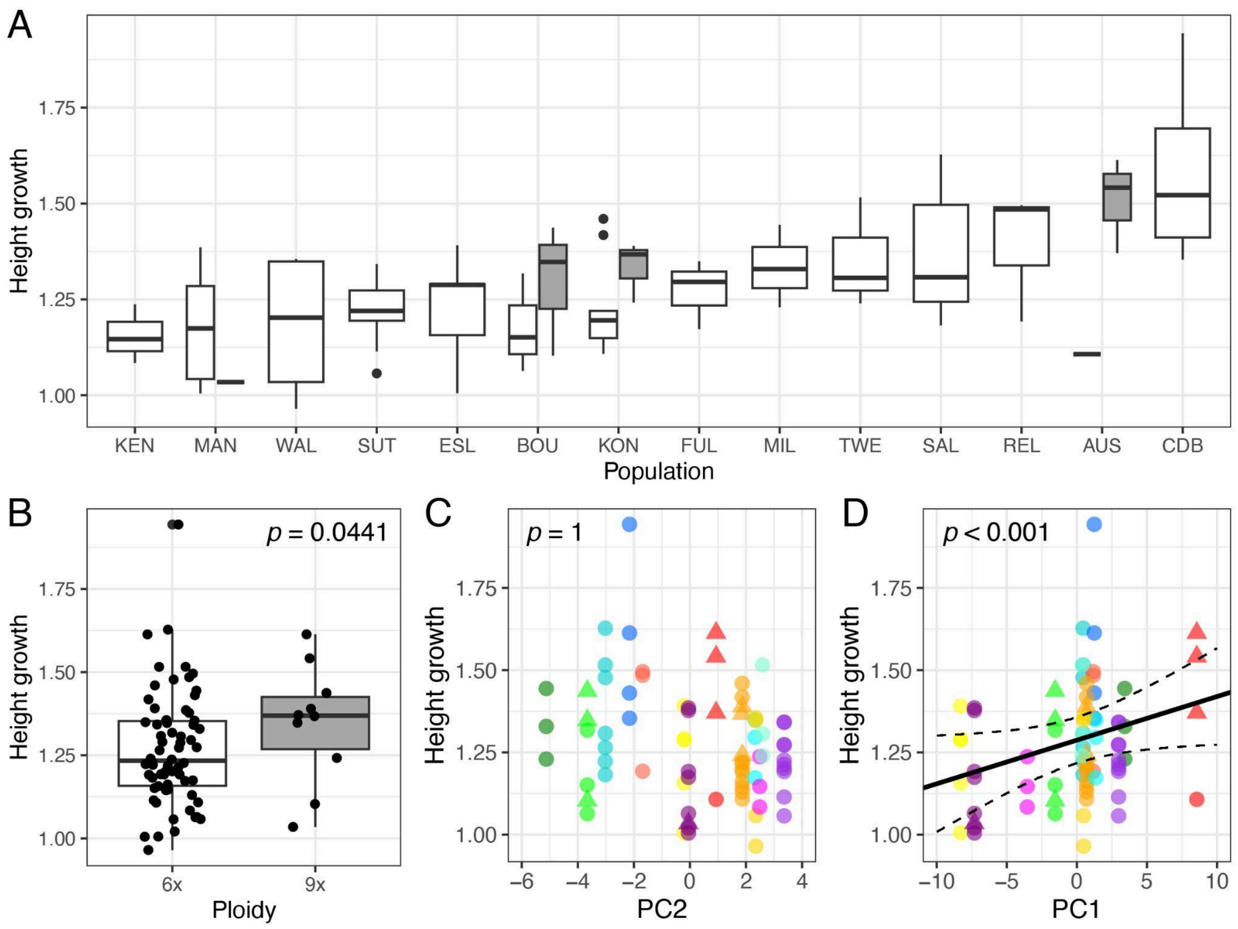


**Figure S23. Variation in the change in plant height (height growth).** See Fig. S5 for figure descriptions.

#### 2.2.3 Number of tillers

We found ploidy (Model 1) and environmental PC1 (Model 2) did not affect the number of tillers (Fig. S8C-D). Environmental PC2 did have a significant effect (Model 2, Fig.

S8E), where the number of tillers increased with precipitation. Having fewer tillers may be beneficial in drier climates, where a conservative growth strategy is needed to prevent water loss by decreasing leaf number (12). We also found the number of tillers to be negatively genetically correlated with basal growth, consistent with previous studies demonstrating a trade-off between yearly aboveground and belowground growth (Fig. 2C, 19).

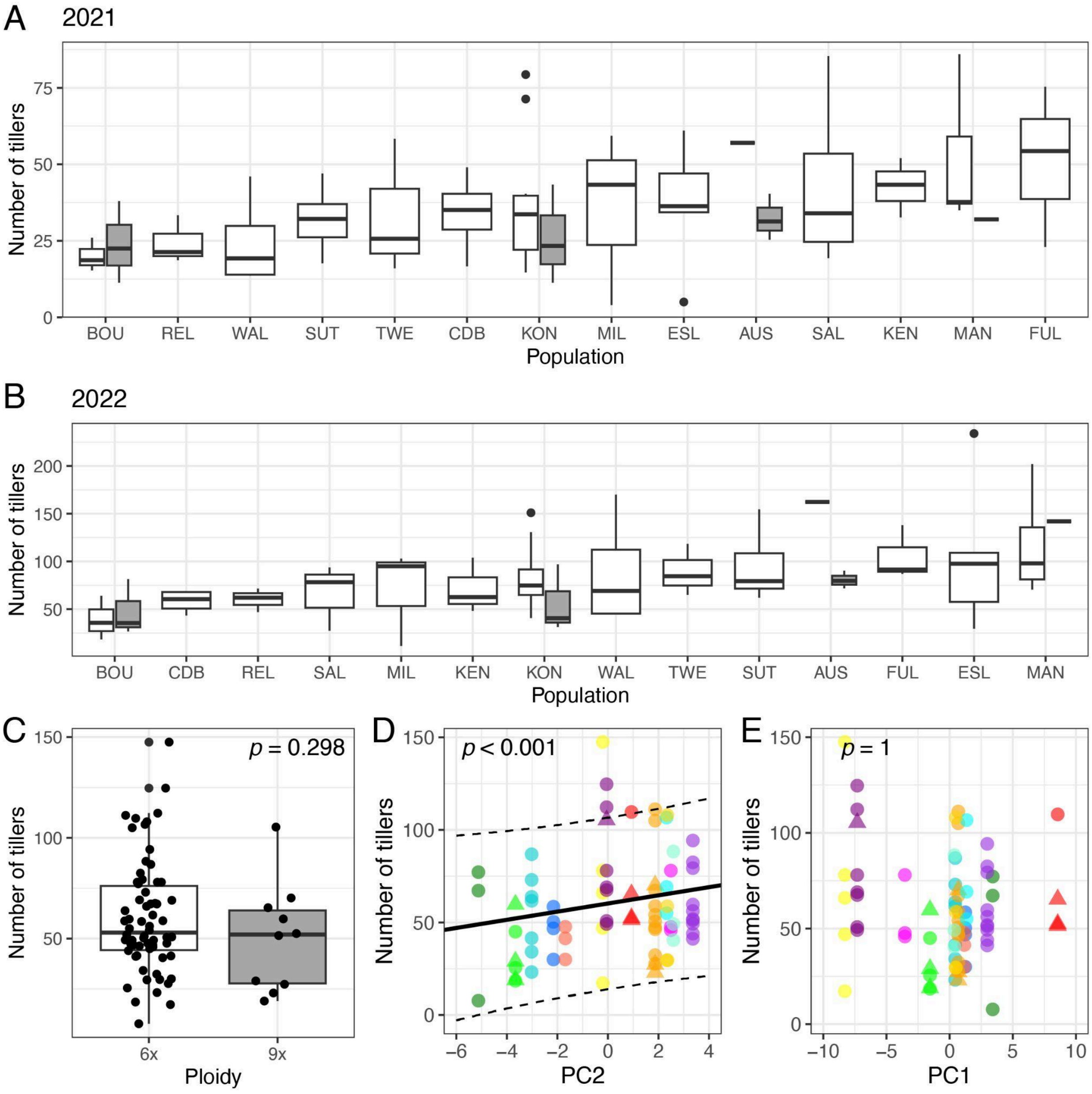


**Figure S24. Variation in the number of tillers.** See Fig. S1 for figure descriptions.

#### 2.2.4 Change in number of tillers (tiller growth)

We found the change in number of tillers (also referred to as tiller growth) was not significantly affected by ploidy (Model 1, Fig. S9B) nor environmental PC1 (Model 2, Fig. S9D). Rather, it was significantly affected by environmental PC2 (Model 2, Fig. S9C), where tiller growth increased with precipitation. Tiller growth was negatively genetically correlated with the number of tillers in both years (Fig. 3C), where plants with the greatest change in tiller number had the fewest tillers. Although the number of tillers was negatively genetically correlated with basal growth, the change in the number of tillers is positively correlated (Fig. 2C). We hypothesize this is because there is a yearly trade-off in above- and belowground biomass but rhizome growth inevitably leads to greater tiller production.

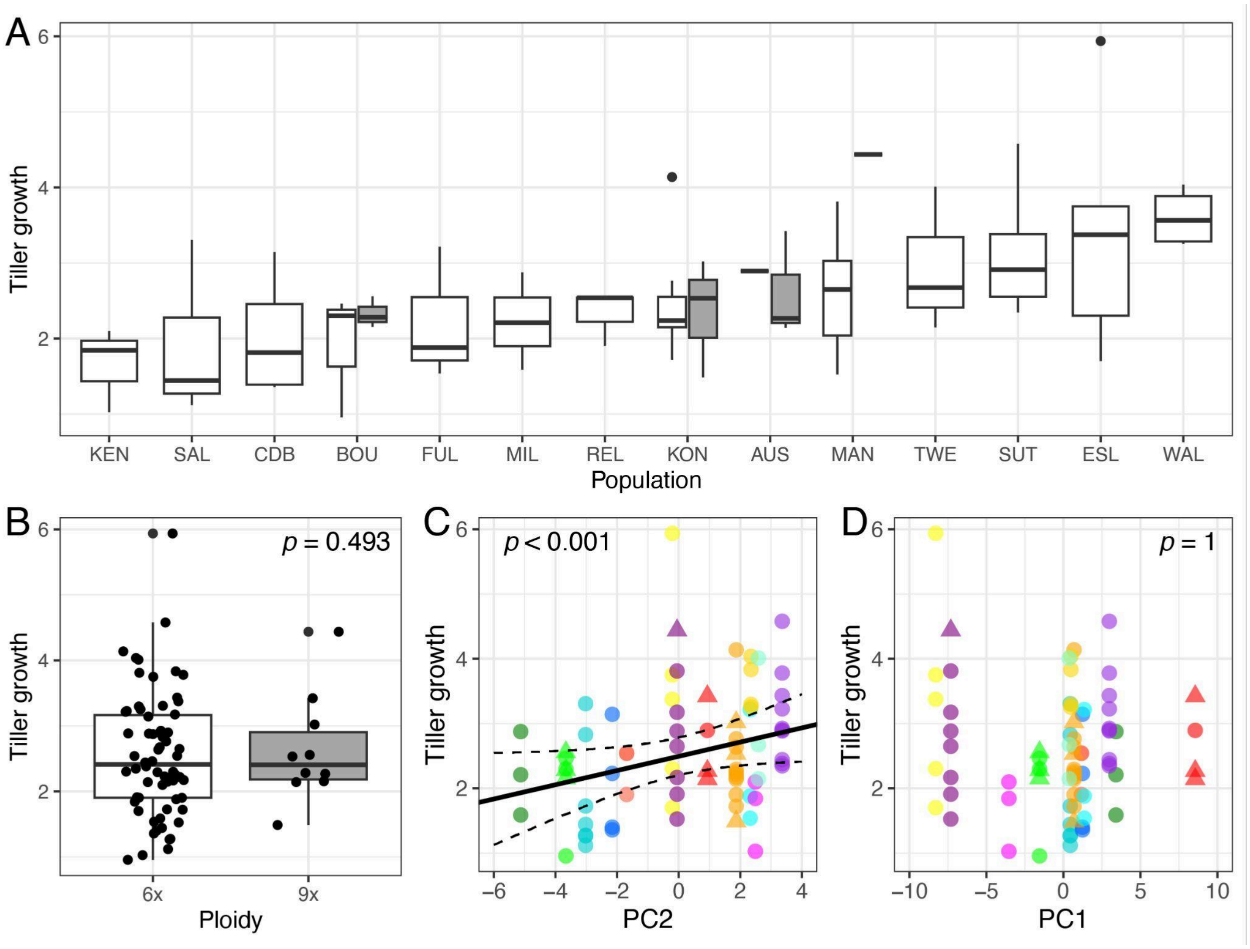


**Figure S25. Variation in the change in number of tillers (tiller growth).** See Fig. S5 for figure descriptions.

#### 2.2.5 Basal growth

Basal growth represents the belowground (rhizome) growth and expansion across the measured years. We found basal growth was not significantly affected by ploidy (Model

1, Fig. S10B) or environmental PC1 (Model 2, Fig. S10D) but was affected by environmental PC2 (Model 2, Fig. S10C). Basal growth significantly increases with precipitation.

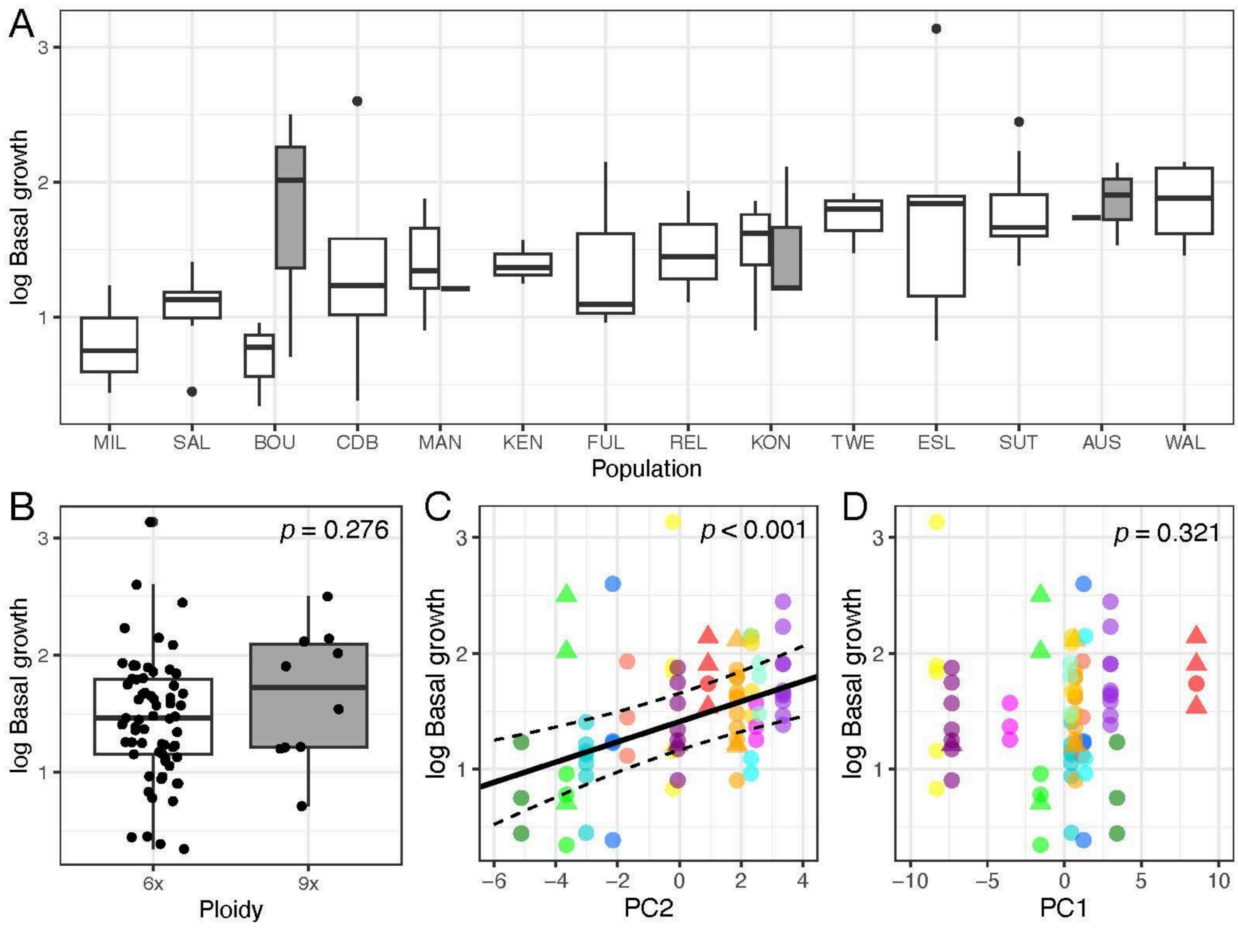


**Figure S26. Variation in basal growth.** See Fig. S5 for figure descriptions. Basal growth was transformed with a log transformation.

#### 2.2.6 Percent of tillers flowering

We found the percent of flowering tillers was not significantly affected by any of the tested variables (Fig. S11). All plants reached flowering before the end of each season.

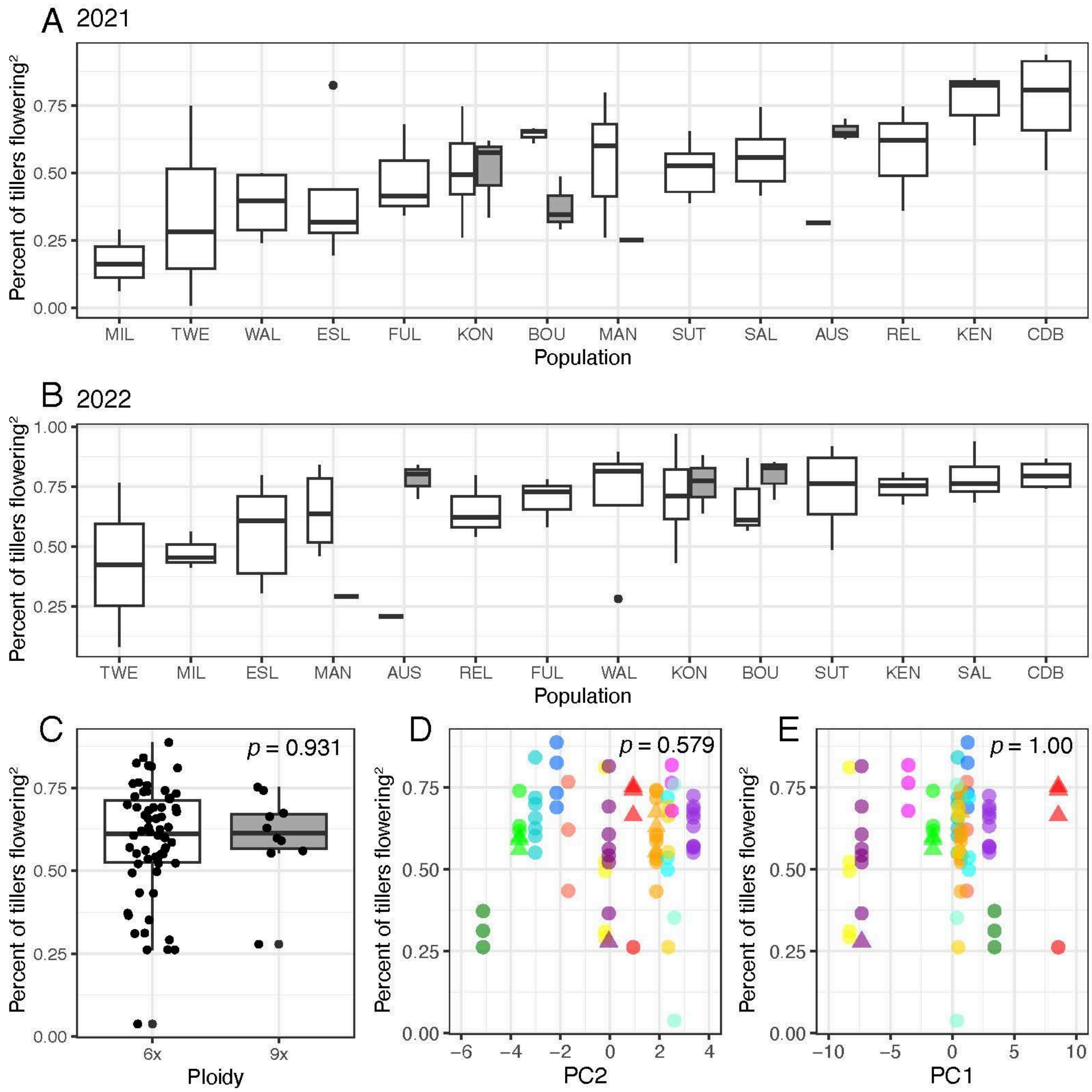


**Figure S27. Variation in the percent of flowering tillers.** See Fig. S1 for figure descriptions. An exponential transformation was applied to the percent of flowering tillers.

#### 2.2.7 Relative growth

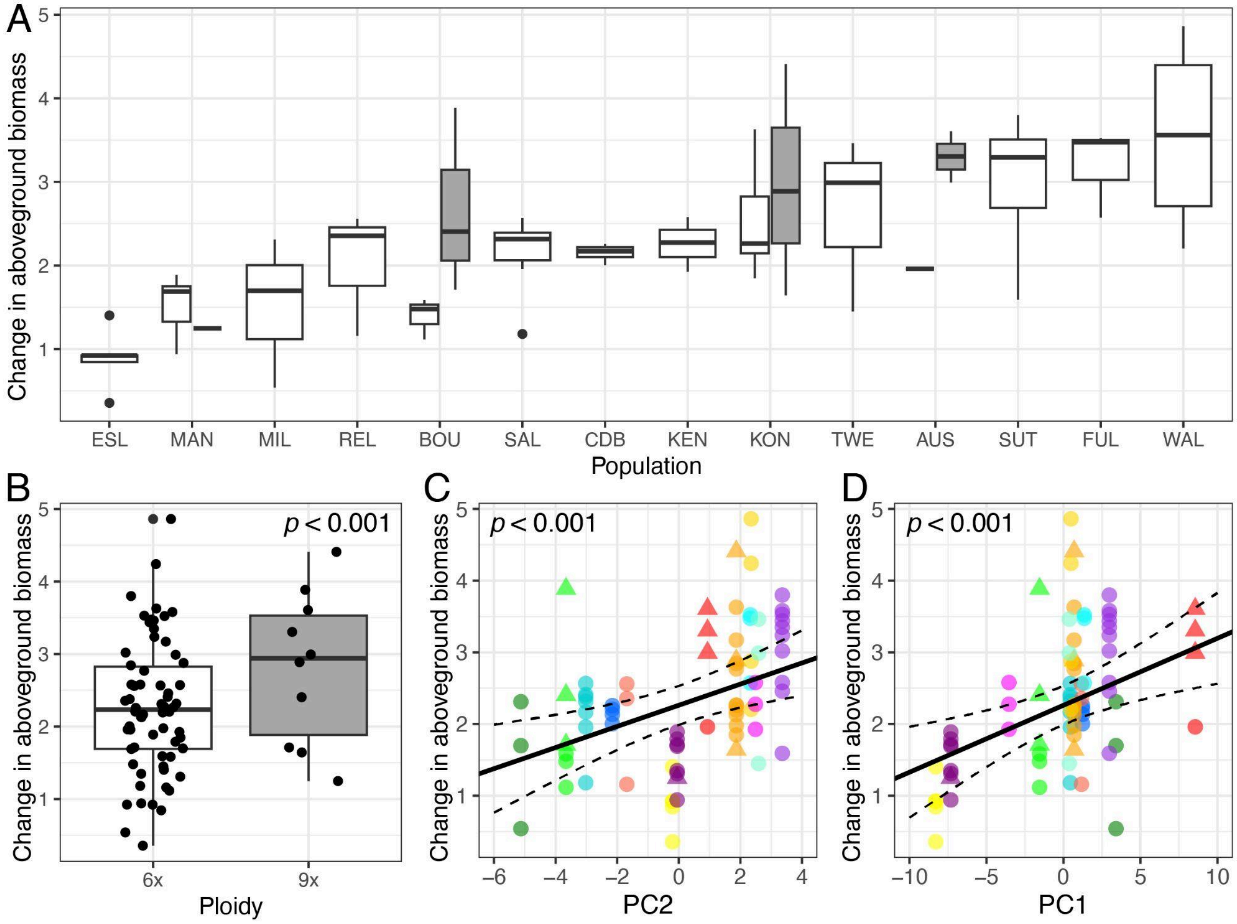


**Figure S28. Variation in relative growth.** See Fig. S5 for figure descriptions. Change in aboveground biomass was measured as the aboveground biomass in year 2 divided year 1.

### 2.3. Stomatal traits

#### 2.3.1 Stomata length

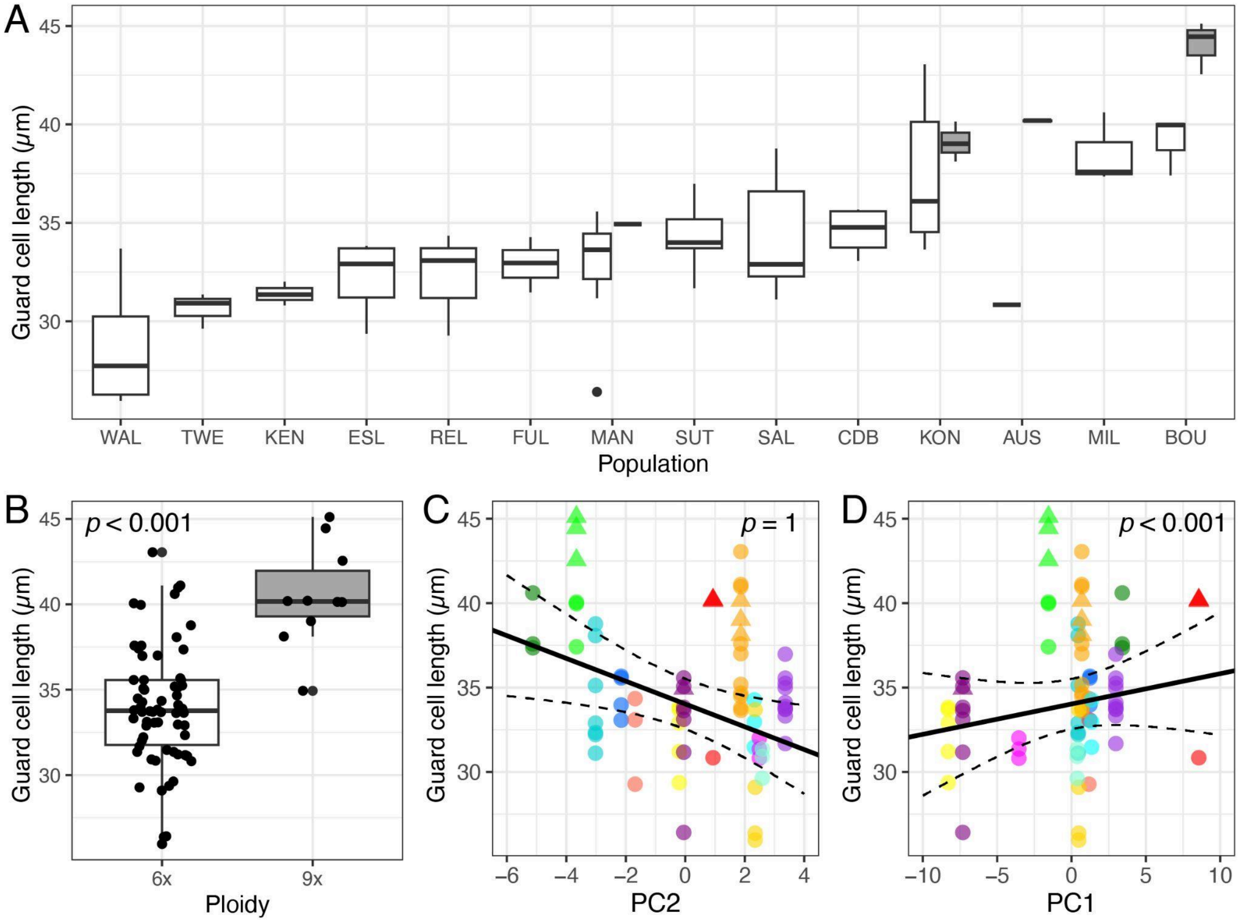


**Figure S29. Variation in stomata length.** See Fig. S5 for figure descriptions.

#### 2.3.2 Stomatal density

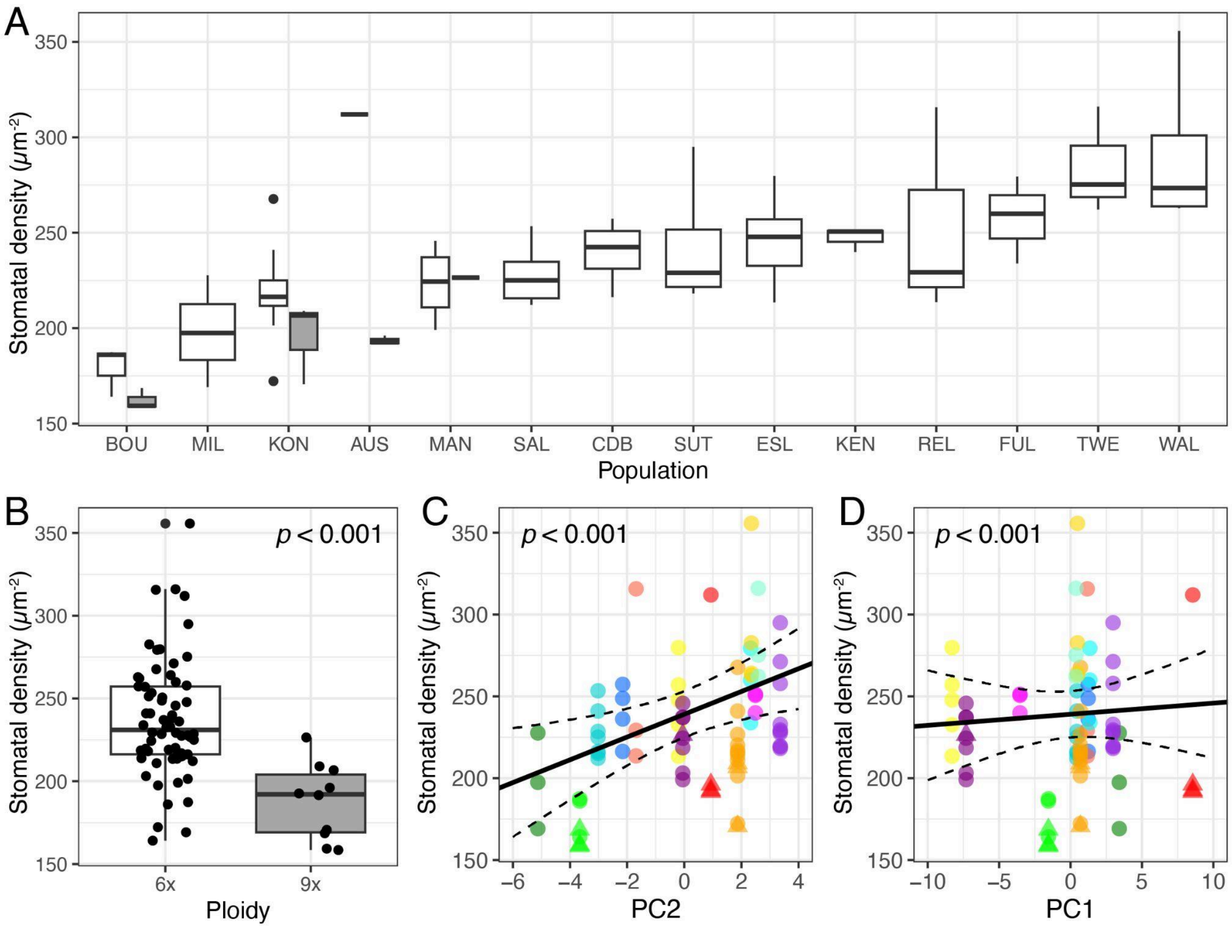


**Figure S30. Variation in stomata density.** See Fig. S5 for figure descriptions.

#### 2.3.1 Stomatal pore index

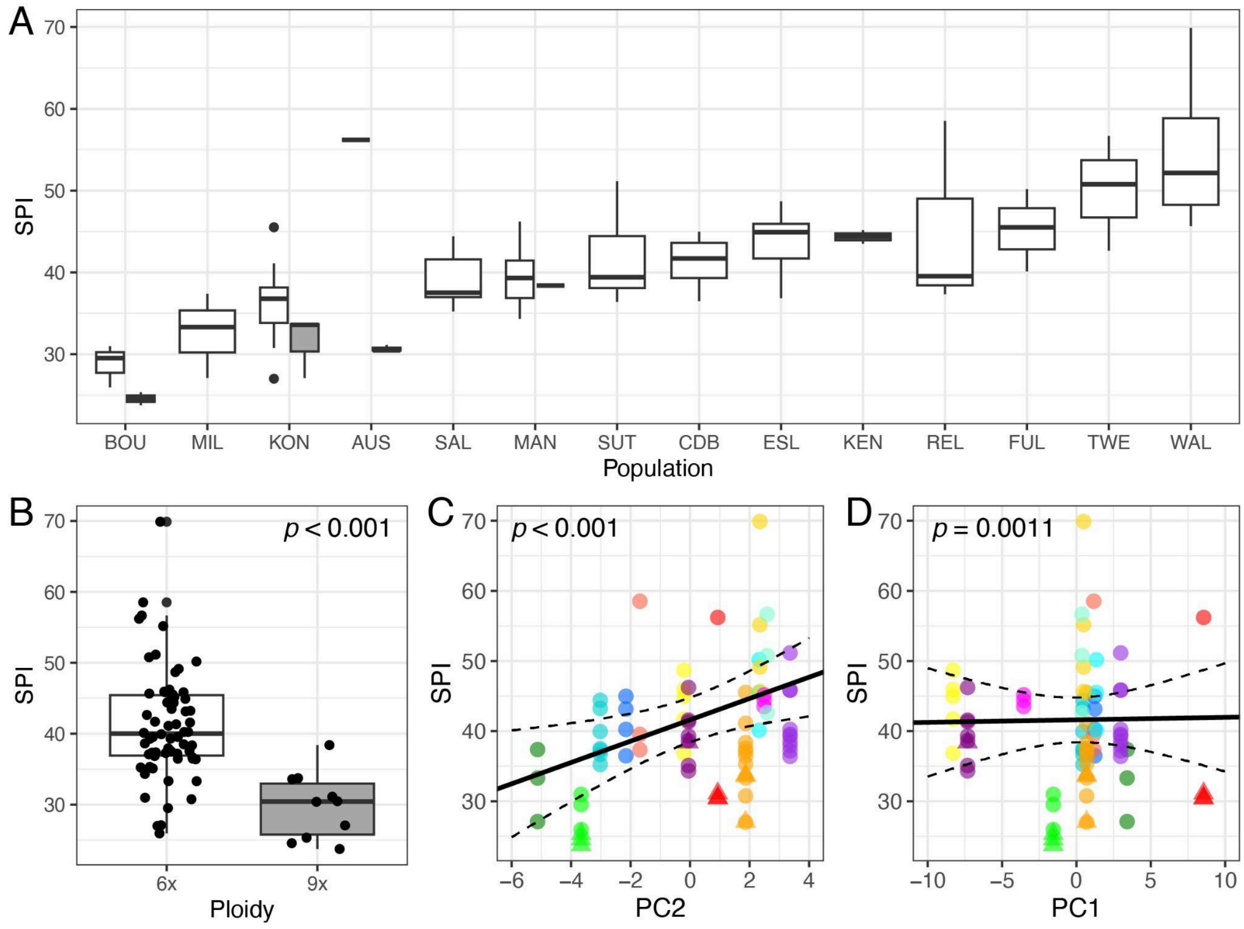


**Figure S31. Variation in stomatal pore index (SPI).** See Fig. S5 for figure descriptions. SPI is unitless.
